# NMR chemical shift assignments of RNA oligonucleotides to expand the RNA chemical shift database

**DOI:** 10.1101/2021.05.20.444957

**Authors:** Yaping Liu, Anita Kotar, Tracy L. Hodges, Kyrillos Abdallah, Mallak H. Taleb, Brayden A. Bitterman, Sara Jaime, Kyle J. Schaubroeck, Ethan Mathew, Nicholas W. Morgenstern, Anthony Lohmeier, Jordan L. Page, Matt Ratanapanichkich, Grace Arhin, Breanna L. Johnson, Stanislav Cherepanov, Stephen C. Moss, Gisselle Zuniga, Nicholas J. Tilson, Zoe C. Yeoh, Bruce A. Johnson, Sarah C. Keane

## Abstract

RNAs play myriad functional and regulatory roles in the cell. Despite their significance, three-dimensional structure elucidation of RNA molecules lags significantly behind that of proteins. NMR-based studies are often rate-limited by the assignment of chemical shifts. Automation of the chemical shift assignment process can greatly facilitate structural studies, however, accurate chemical shift predictions rely on a robust and complete chemical shift database for training. We searched the Biological Magnetic Resonance Data Bank (BMRB) to identify sequences that had no (or limited) chemical shift information. Here, we report the chemical shift assignments for 12 RNA hairpins designed specifically to help populate the BMRB.

## Biological context

RNA is a multifaceted biomolecule that plays important functional and regulatory roles in the cell. In the simplest case, RNA has a crucial role in the central dogma of molecular biology. Not only do messenger (m) RNAs serve as the template for protein synthesis in the cell, the cellular machinery responsible for protein synthesis, the ribosome, is largely composed of RNA molecules. Outside of its role in the central dogma, RNAs contribute to the overall health of the cell by controlling splicing (Papasaikas and Valcarcel, 2016), translation efficiency (Rodnina, 2016), and genomic stability (Theimer and Feigon, 2006). More recently, RNA molecules have been used as the foundation for vaccines (Corbett et al., 2020; Pardi et al., 2018), investigated as therapeutics (Burnett and Rossi, 2012), and have been identified as potential therapeutic targets themselves (Howe et al., 2015; Meyer et al., 2020; Warner et al., 2018).

Despite the importance of RNAs in biology, structural elucidation of RNA molecules lags significantly behind their protein counterparts. As of May 2021, the Protein Data Bank (PDB) (Berman et al., 2000a; Berman et al., 2000b) has 173,534 protein-containing structures and only 5,364 RNA-containing structures. Similarly, the Biological Magnetic Resonance Data Bank [BMRB (Ulrich et al., 2008)] contains 13,420 protein depositions and only 467 RNA depositions. This imbalance in our RNA structural knowledge has led to a similar disparity in our mechanistic understandings of the functions of these RNAs.

Chemical shifts are a fundamental indicator of molecular structure and are at the heart of all NMR studies (Ebrahimi et al., 2001; Fares et al., 2007; Ghose et al., 1994; Giessner-Prettre and Pullman, 1987; Rossi and Harbison, 2001; Sripakdeevong et al., 2014). However, assignment of chemical shifts remains a laborious and often rate-limiting step in the determination of biomolecular structure and characterization of biomolecular dynamics and ligand interactions. A number of groups are actively working to develop and improve methods to predict chemical shifts of nuclei in proteins (Han et al., 2011; Meiler, 2003), DNA (Kwok and Lam, 2013; Lam, 2007; Lam et al., 2007; Ng and Lam, 2015), and RNAs (Aeschbacher et al., 2013; Bahrami et al., 2012; Barton et al., 2013; Brown et al., 2015; Frank et al., 2013; Frank et al., 2014; Krahenbuhl et al., 2014; Wang et al., 2021). In our experience, a good prediction can significantly reduce the time required for the assignment of chemical shifts. However, the accuracy of these algorithms requires a robust database of chemical shifts that represent a wide variety of both sequence and structural motifs. At the beginning of this study, we found only 244 entries at the BMRB for RNA molecules that were usable with our chemical shift analysis tools. The sparseness of this database limits the accuracy of the available tools.

In a canonical A-form helix, the neighboring base pairs have the most significant impact on the chemical shifts of the central base pair (Barton et al., 2013). There are 64 possible canonical base pair triplets, which we have defined as a group. Because the specific identity of the base pairs flanking this core sequence has a lesser effect on the chemical shift of the central nucleotide, we considered nucleotide type (purine or pyrimidine) rather than nucleotide identity. This is consistent with the attributes used in the current prediction algorithm used in NMRFx Analyst (Marchant et al., 2019). If each of these “groups” is flanked by a purine-pyrimidine base pair or a pyrimidine-purine base pair at the 5′- and 3′-ends, there are 256 possible combinations of 5 base pair sequences. With these 256 sequences in hand, we examined the BMRB for aromatic protons and carbon chemical shifts and identified four “groups” of RNAs whose chemical shifts were underrepresented (**Fig. 1**). We prepared 12 unique RNAs, three from each group, for chemical shift assignment.

**Figure 1.**
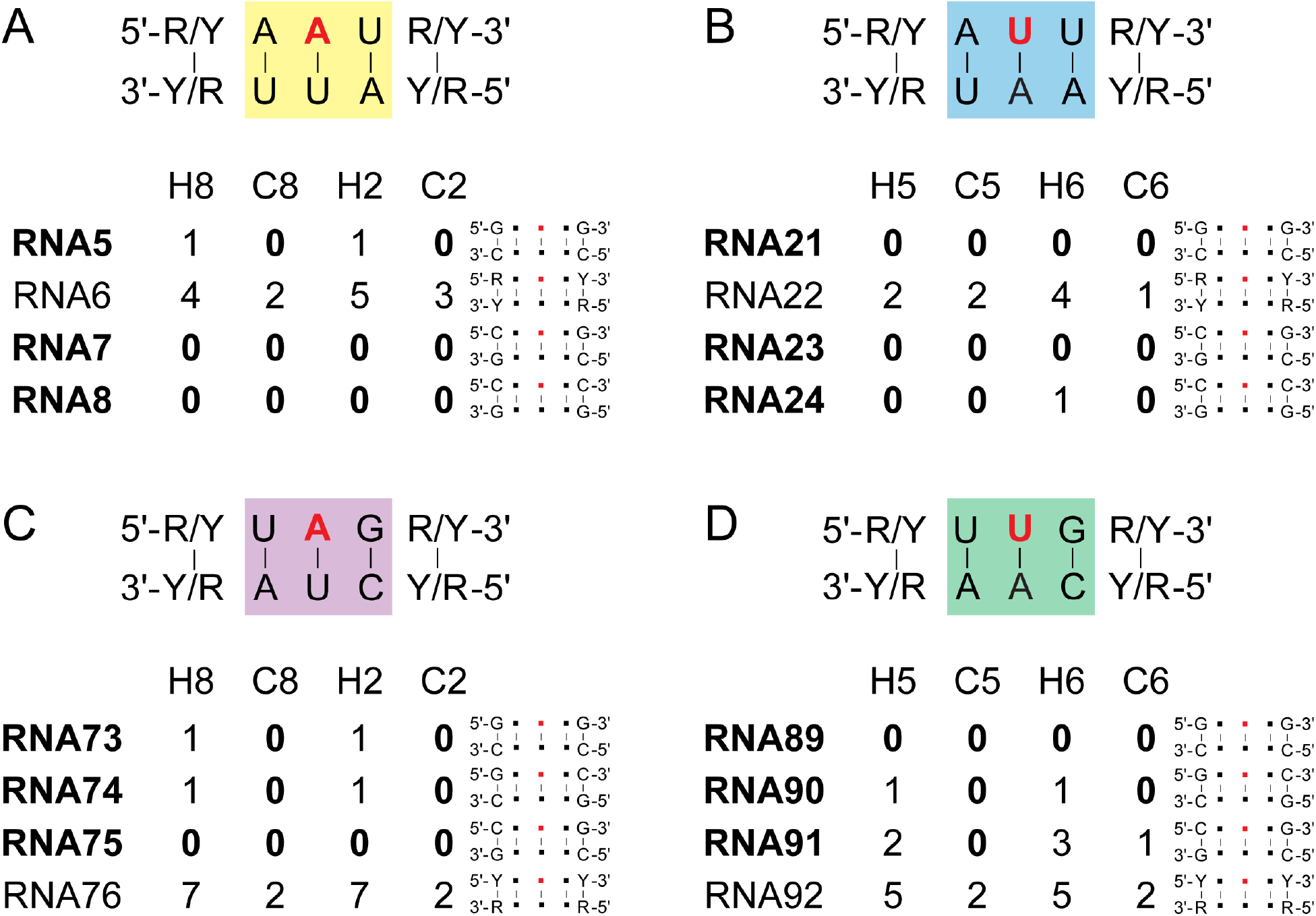
Overview of RNA sequences with missing assignments. Four groups of RNAs were identified A) Group 1, RNAs 5-8, B) Group 2, RNAs 21-24, C) Group 3, RNAs 73-76, D) Group 4, RNAs 89-92. Red nucleotide indicates the nucleotide with missing chemical shifts. The common triple of base pairs for each group is highlighted in a colored box. Sequence variations at flanking nucleotides were examined. The occurrence of aromatic proton and carbon chemical shifts in the BMRB for each RNA are indicated. Flanking sequences for each RNA are shown. R, purine; Y, pyrimidine.

The RNAs examined in this study were designed to fill specific gaps in the BMRB chemical shift database. Additionally, these RNAs represent helical regions from a diverse set of biological RNAs including ribosomes, tRNAs, riboswitches, internal ribosome entry sites, and viral RNAs (**Table S1**) as identified via the RNA fragments search engine and database [RNA FRABASE, (Popenda et al., 2008; Popenda et al., 2010)]. The RNA FRABASE searches for RNA fragments based on known structures, therefore there are likely many more biologically-relevant RNAs that are represented by these RNA oligonucleotides. These additional chemical shift assignments, which are a step towards a more complete chemical shift database, will benefit the RNA NMR community.

## Materials and Methods

### Identification of RNA sequences that are underrepresented in the BMRB

The identification of RNA sequences that are underrepresented in the BMRB was done using the tools previously developed for training models for the prediction of RNA chemical shifts (Brown et al., 2015). We used these tools to analyze NMR-STAR files containing RNA data that we downloaded from the BMRB. The downloaded NMR-STAR files were analyzed with the scripts developed for the prediction model training. This process automatically identifies PDB files to generate secondary structures from and does various validation checks. After this process was complete, 244 files were available for the subsequent analysis. This analysis generated a table of chemical shifts and attributes as used in the prediction software. The tabular data was searched for all 256 five-residue sequences described above and a new table generated containing a line for each of the 256 sequences. Each line contains the number of measured chemical shifts for each of the aromatic proton and carbon atoms for the corresponding sequence. Of these 256 sequences, 182 of them have at least one measurement of all the aromatic proton and carbons (**Table S2**). The remaining 74 entries were missing the measurement of the chemical shift of at least one atom.

### Construct design and template preparation

DNA oligonucleotides corresponding to the 12 RNA constructs (**Table 1**) were ordered (Integrated DNA Technologies) with 2′-*O*-methoxy modifications at the two 5′-most positions to reduce non-templated transcription (Kao et al., 1999). To generate the templates for transcription, DNA oligonucleotides (**Table 1**) were annealed with an oligo corresponding to the T7 promoter sequence (5′-TAATACGACTCACTATA-3′). Briefly, 40 µL of each DNA oligonucleotide (200 µM) was individually mixed with 20 µL of the T7 promoter sequence oligonucleotide (600 µM). These samples were placed in boiling water for 3 minutes and left in the water bath to slowly cool to room temperature over at least 2 hours. The annealed template was then diluted with 940 µL H_2_O to produce the partially double-stranded DNA templates for transcription.

**Table 1.**
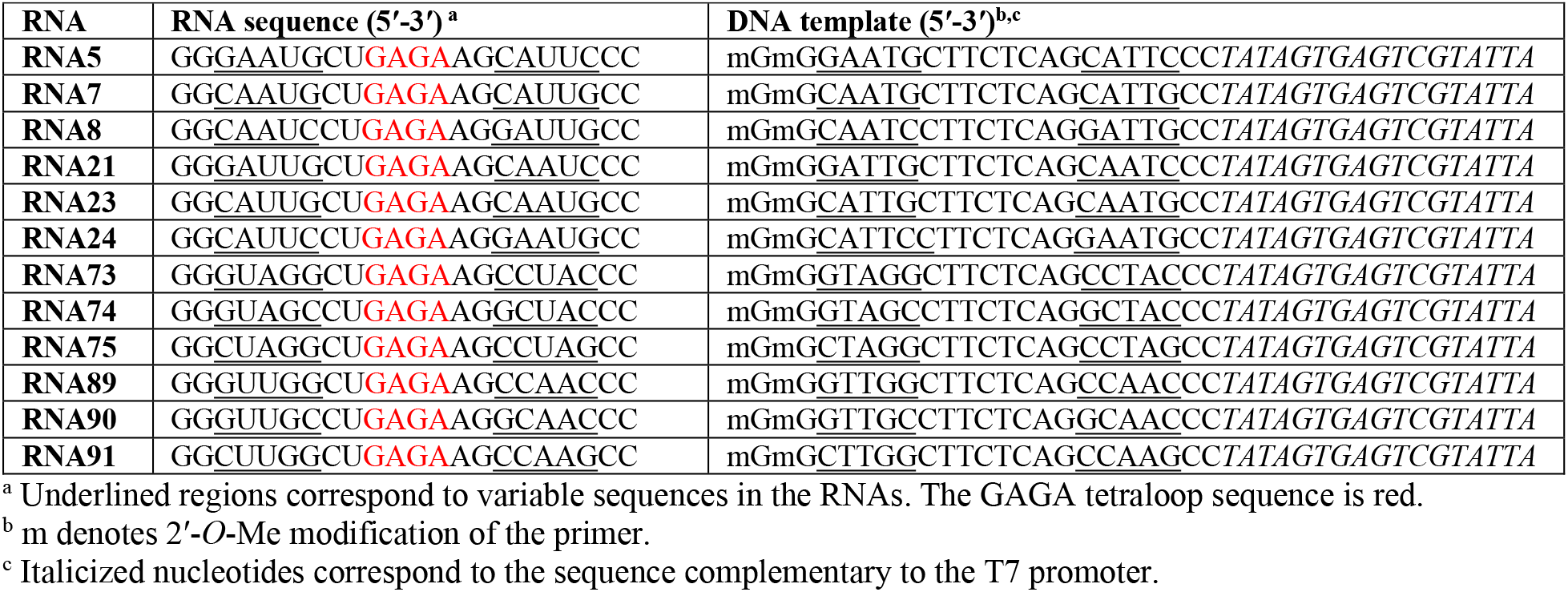
RNA constructs.

### RNA preparation

RNA transcription conditions were optimized prior to large scale preparation. RNAs were prepared by *in vitro* transcription in transcription buffer (40 mM Tris-base, 5 mM DTT, 1 mM Spermidine, 0.01% Triton-X, pH 8.5) with addition of 2-6 mM NTPs, 10-20 mM MgCl_2_, 8-13 pmol annealed DNA template, 0.2 U/mL yeast inorganic pyrophosphatase (New England Biolabs) (Cunningham and Ofengand, 1990), ∼15 µM T7 RNA polymerase, and 10-20% (vol/vol) DMSO (Helmling et al., 2015). Reactions were incubated at 37 °C for 3-4 hours and then quenched using a solution of 7 M urea and 250 mM EDTA (pH 8.5). The quenched transcription mixture was loaded onto 16% preparative-scale denaturing gels for RNA purification. The RNA band was identified by UV shadowing, excised, and electroeluted from the gel for ∼24 h using a membrane trap elution system (Elutrap, Whattman). RNAs were spin concentrated, washed with 2 M high-purity sodium chloride, and exchanged into water using Amicon Centrifugal Filter Units (Millipore, Sigma). RNA purity was checked by running RNA on a 16% analytical denaturing gel.

### NMR spectroscopy

RNA samples (300-400 µM in 300 µL) were prepared in 50 mM potassium phosphate buffer, pH = 7.4. Samples were lyophilized and rehydrated in an equal volume of D_2_O (99.8%, Cambridge Isotope Laboratories, Inc.). 2D ^1^H-^1^H NOESY, ^1^H-^1^H TOCSY, and ^1^H-^13^C HMQC spectra were recorded for each RNA at 25 °C (see **Table 2** for experimental parameters). NMR spectra were collected on a 600 MHz Bruker AVANCE NEO spectrometer equipped with a 5 mm TCI cryogenic probe (University of Michigan BioNMR Core). NMR data were processed with NMRFx (Norris et al., 2016) and NMRPipe (Delaglio et al., 1995) and analyzed with NMRViewJ (Johnson and Blevins, 1994). ^1^H chemical shifts were referenced to water and ^13^C chemical shifts were indirectly referenced from the ^1^H chemical shift (Wishart et al., 1995).

**Table 2.**
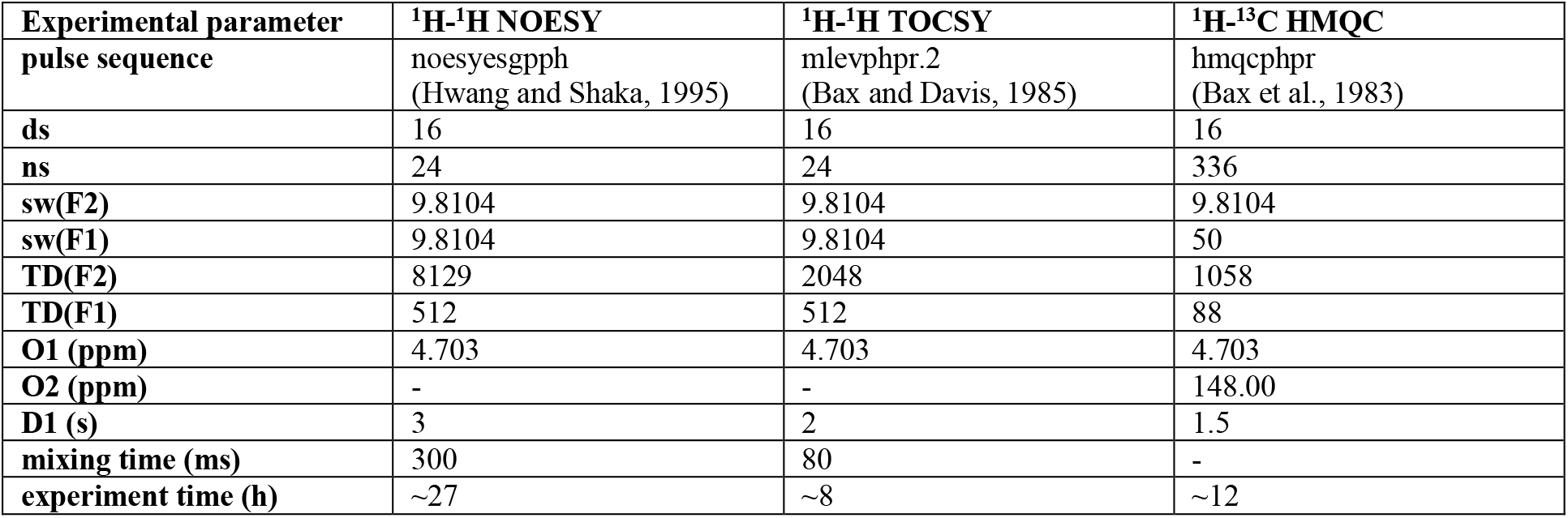
Experimental parameters for NMR data acquisition.

### Assignments of helical regions

For all the RNA oligos, the nonexchangeable proton assignments were unambiguously assigned through using 2D ^1^H-^1^H NOESY (Hwang and Shaka, 1995), 2D ^1^H-^1^H TOCSY (Bax and Davis, 1985), and ^1^H-^13^C HMQC (Bax et al., 1983) experiments. The 2D ^1^H-^1^H TOCSY spectrum yields strong H5-H6 cross-peaks, reporting on the number of pyrimidines (cytosine and uracil) in an RNA molecule, and was used to identify the H5-H6 correlations in the corresponding ^1^H-^1^H NOESY spectrum. The ^1^H-^1^H NOESY spectrum was used to make assignments of sequential nucleotides in A-helical regions. Near the diagonal, sequential aromatic-aromatic proton correlations were observable for many regions of each RNA. In the aromatic-anomeric region, sequential assignments were possible by following the NOE pattern for A-helical regions. Briefly, the H6/H8 of each residue (i) has NOE with its own H1′ and the H1′ of the preceding nucleotide (i-1). Assignments could be confirmed by examining additional NOEs, for example the aromatic H6/H8 proton of a residue (i) has a weak, but detectable NOE to the H5 of a following (i+1) pyrimidine. Additionally, the position of the adenosine C2 proton (H2) in the interior of the helix provides rich NOE data informing on both intra-strand (sequential) and cross-strand connectivities. When part of an A-form helix, the adenosine H2 (i) has NOEs to its own H1′ (i), to the H1′ of its 3′ residue (i+1), and an inter-strand cross-peak to H1′ of the residue (j+1) 3′ of the base to which it is paired (j). The 2D ^1^H-^13^C HMQC was used to help distinguish C2-H2 (^13^C δ ∼152 ppm) from C6-H6 and C8-H8 resonances (^13^C δ ∼139 ppm).

### GAGA tetraloop

Hairpin or stem-loop structures are pervasive in RNAs. RNA hairpins with a four nucleotide loop are known as tetraloops and are the most common size of loop (Antao and Tinoco, 1992). All RNA constructs were designed to contain a GAGA tetraloop with a U-A closing base pair (**Fig. 2**). The GAGA tetraloop was chosen due to its structural stability (Dale et al., 2000; Sheehy et al., 2010) and characteristic signals in the ^1^H-^1^H NOESY spectrum (**Fig. 3**). The NOE walk of the GAGA tetraloop presents a unique pattern which was helpful to both confirm proper folding of the RNA and facilitate resonance assignments. The three-dimensional structure of the nucleotides in the GAGA tetraloop cause a dramatic change in the H1′ frequency of the nucleotide following the loop (A14 in these constructs), out of the typical anomeric proton chemical shift range and to a smaller (upfield) chemical shift (Jucker et al., 1996).

**Figure 2.**
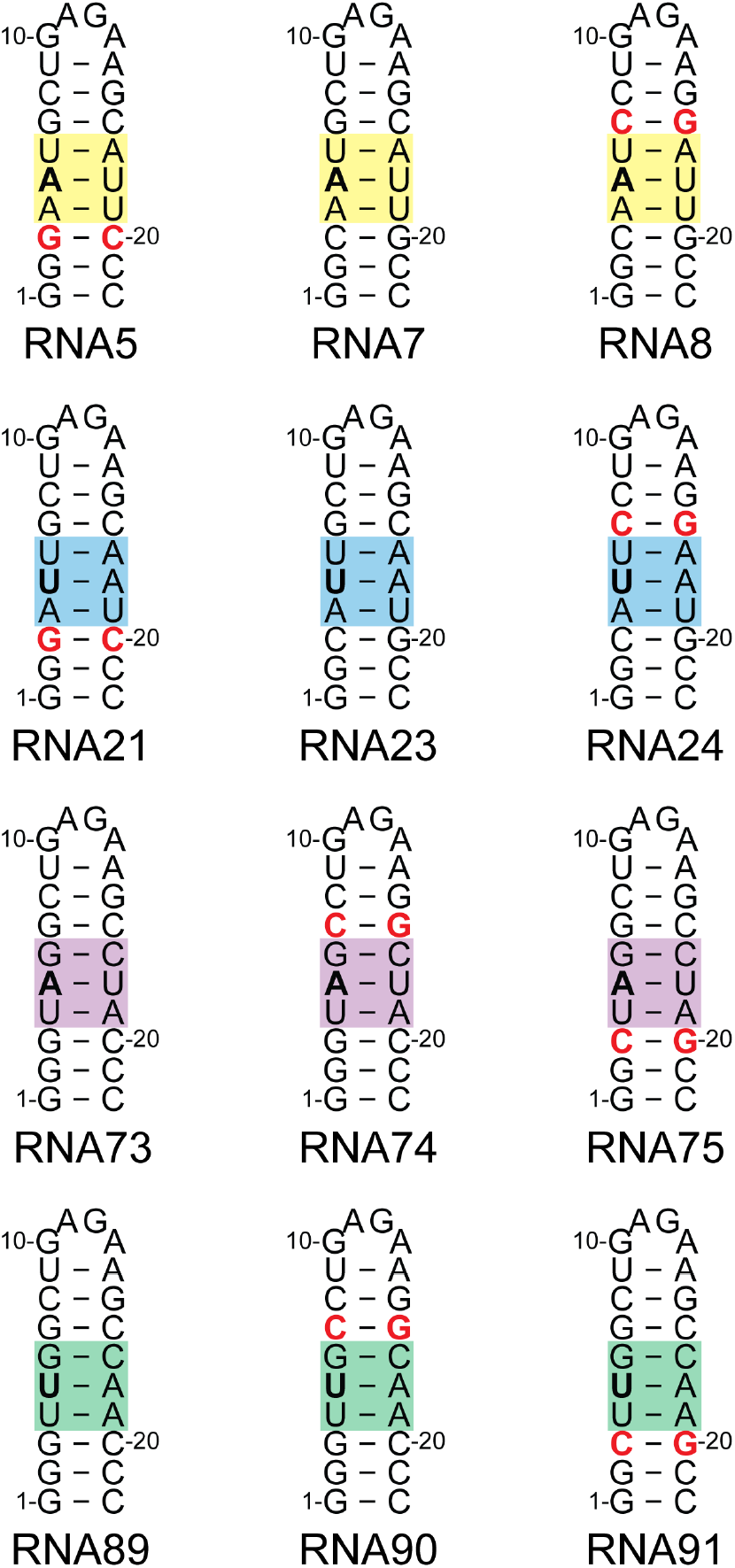
RNA constructs used in this study. Red nucleotides indicate the variable sequence relative to a single example for each group. The common triple of base pairs for each group is highlighted in a colored box.

**Figure 3.**
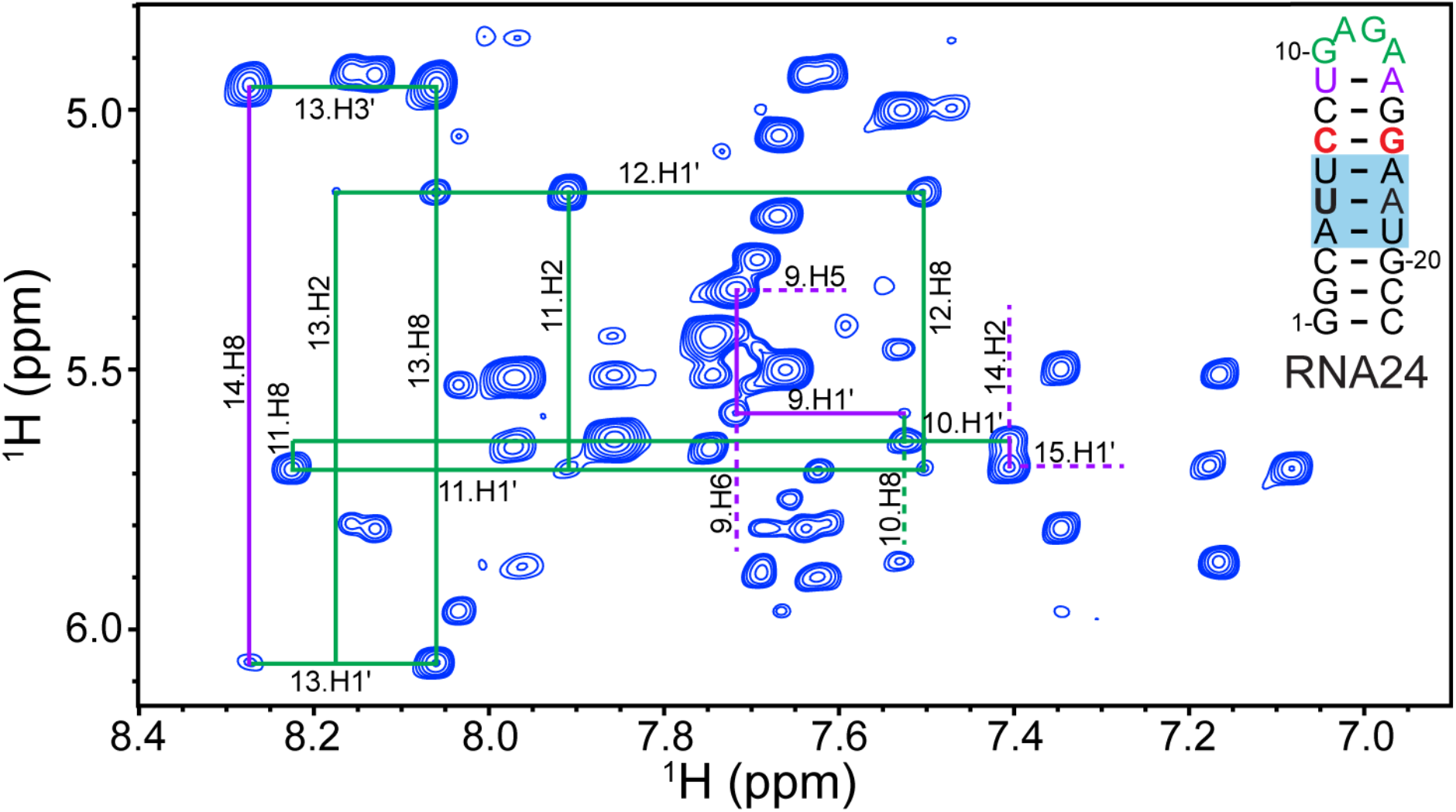
^1^H-^1^H NOESY spectrum of RNA24, highlighting the chemical shift assignments for the GAGA tetraloop. Tetraloop resonances are connected with green lines, resonances from flanking sequences are connected with purple lines. Dashed lines are used in crowded regions to indicate resonance assignment.

### Assignments and data deposition

Assigned ^1^H-^1^H NOESY and ^1^H-^13^C HMQC spectra for the 12 RNAs are presented in **Figs. 4-7** and **Supplemental Figs. S1-S8**. In this work, we have unambiguously assigned all 330 aromatic C6-H6, C8-H8, and C2-H2 correlations. For the 66 adenosine residues, we have assigned 100% of the aromatic and anomeric protons (A-H2, A-H8, and A-H1′). We were able to assign 86% of the adenosine H2′ and 74% of adenosine H3′ protons (57/66 and 49/66, respectively). For the 90 guanosine residues, we have assigned 100% of the aromatic and anomeric protons (G-H8, and G-H1′). We were able to assign 89% of the guanosine H2′ and 86% of guanosine H3′ protons (80/90 and 77/90, respectively). For the 66 cytosine residues, we have assigned 100% of the aromatic and anomeric protons (C-H5, C-H6, and C-H1′). We were able to assign 89% of the cytosine H2′ and 77% of cytosine H3′ protons (59/66 and 51/66, respectively). Finally, for the 42 uracil residues, we have assigned 100% of the aromatic and anomeric protons (U-H5, U-H6, and U-H1′). We were able to assign 90% of the uracil H2′ and 90% of uracil H3′ protons (38/42 and 38/42, respectively).

**Figure 4.**
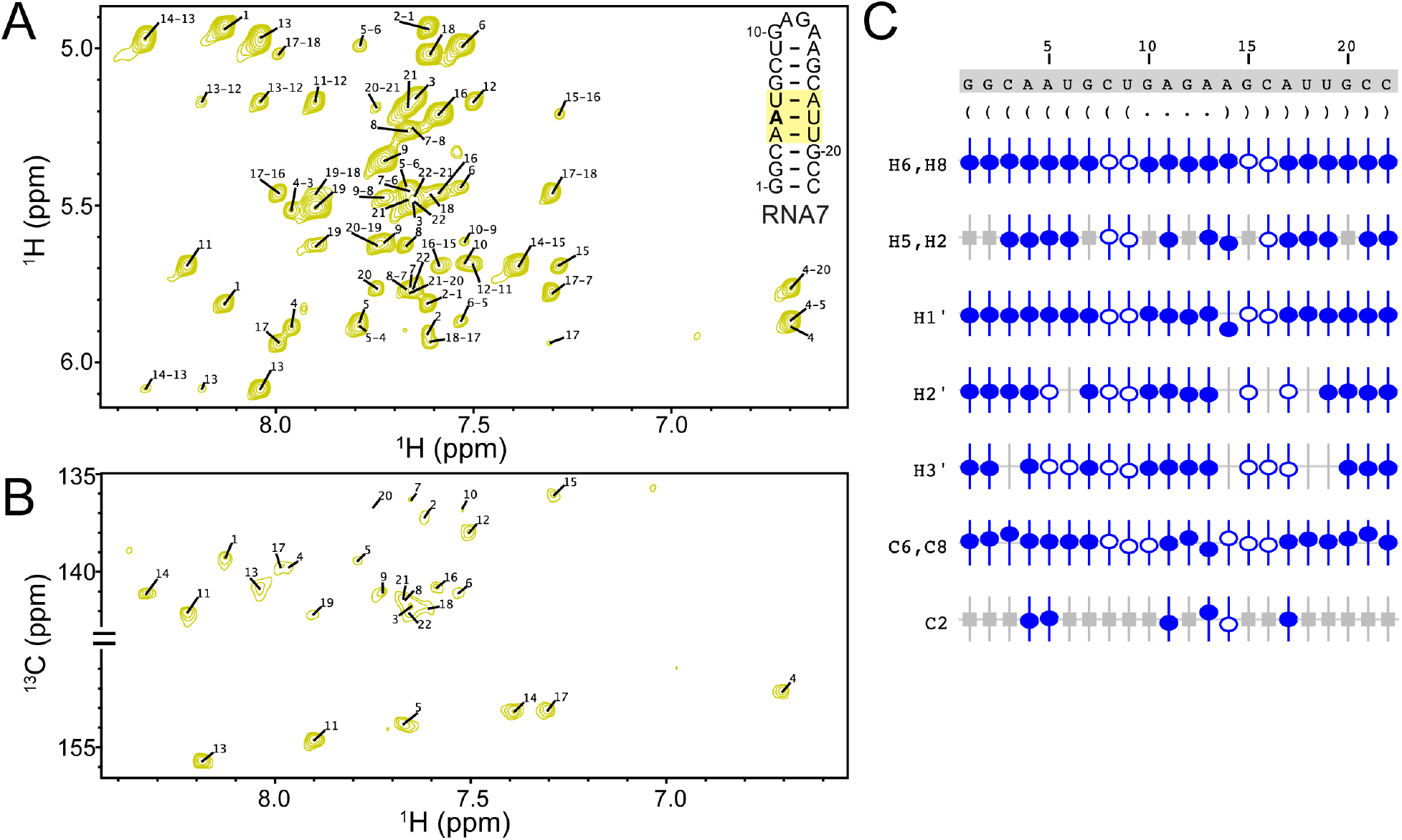
RNA7 chemical shift assignments. A) ^1^H-^1^H NOESY, B) ^1^H-^13^C HMQC, C) Summary of the sequence, secondary structure, and assignment validation for RNA7. The secondary structure is shown in Vienna format. NMRViewJ assignment summary to validate proton (H6/H8, H5/H2, H1′, H2′, H3′) and carbon (C6/C8, C2) assignments (bottom). Assigned chemical shifts for specific atoms are indicated with open and filled circles. The vertical offset of the circles indicates the deviation from the predicted values for that atom. Filled circles indicate that there are chemical shifts for atoms with the same set of attributes in the BMRB. Open circles indicate atoms that have a prediction, but for which no exact matches of the attributes are available in the BMRB. Grey boxes represent atoms that are not present in a given base.

**Figure 5.**
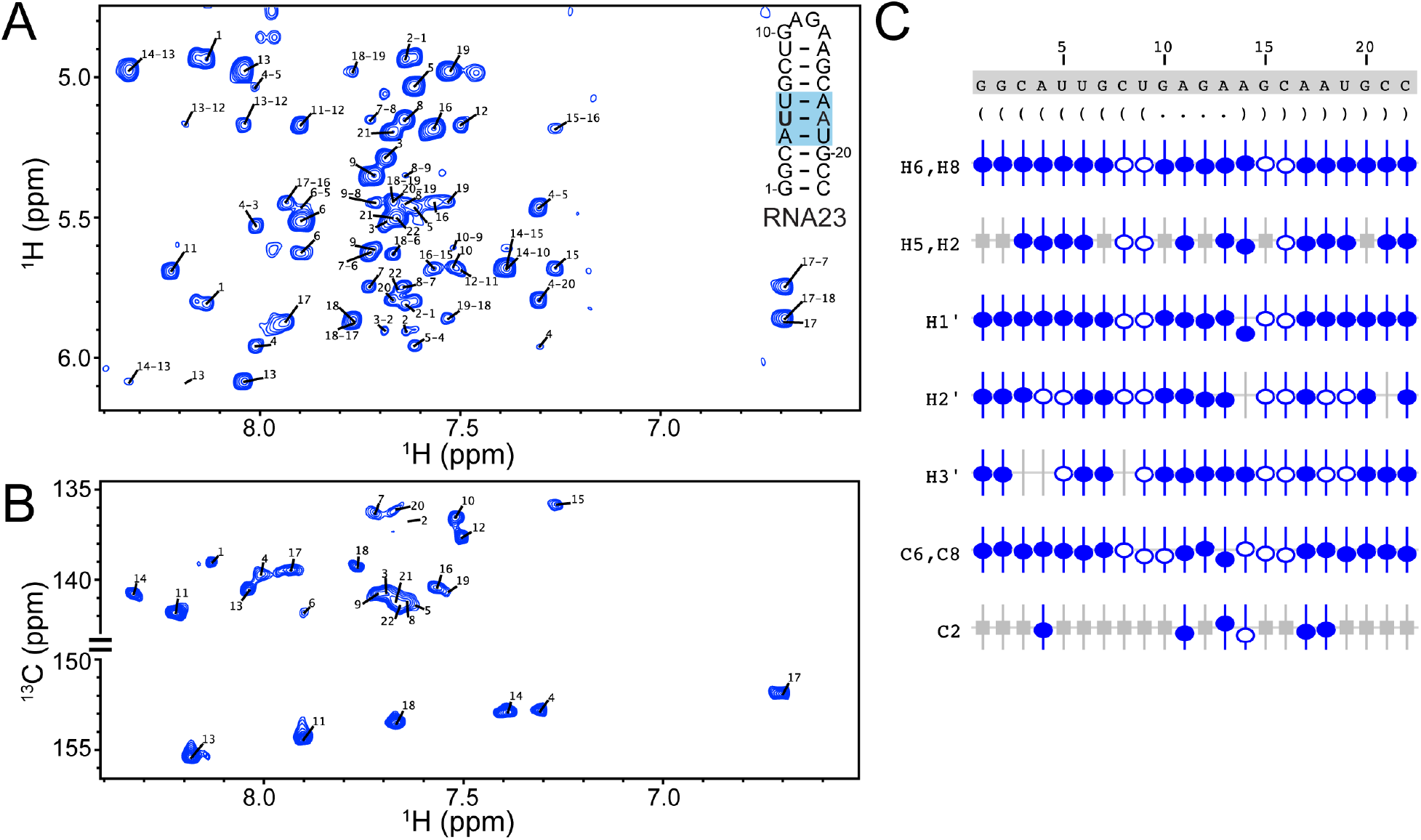
RNA23 chemical shift assignments. A) ^1^H-^1^H NOESY, B) ^1^H-^13^C HMQC, C) Summary of the sequence, secondary structure, and assignment validation for RNA23. The secondary structure is shown in Vienna format. NMRViewJ assignment summary to validate proton (H6/H8, H5/H2, H1′, H2′, H3′) and carbon (C6/C8, C2) assignments (bottom). Assigned chemical shifts for specific atoms are indicated with open and filled circles. The vertical offset of the circles indicates the deviation from the predicted values for that atom. Filled circles indicate that there are chemical shifts for atoms with the same set of attributes in the BMRB. Open circles indicate atoms that have a prediction, but for which no exact matches of the attributes are available in the BMRB. Grey boxes represent atoms that are not present in a given base.

**Figure 6.**
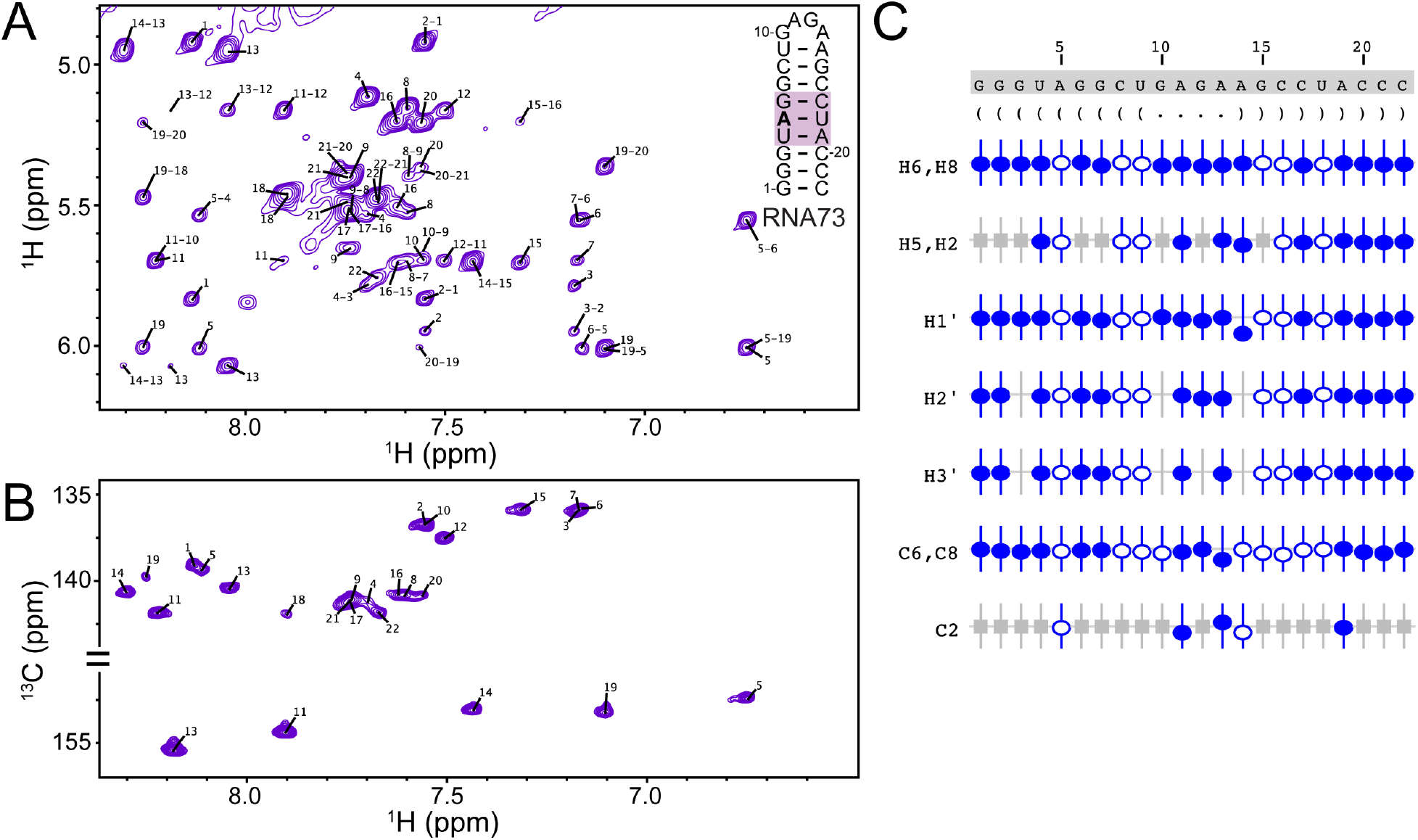
RNA73 chemical shift assignments. A) ^1^H-^1^H NOESY, B) ^1^H-^13^C HMQC, C) Summary of the sequence, secondary structure, and assignment validation for RNA73. The secondary structure is shown in Vienna format. NMRViewJ assignment summary to validate proton (H6/H8, H5/H2, H1′, H2′, H3′) and carbon (C6/C8, C2) assignments (bottom). Assigned chemical shifts for specific atoms are indicated with open and filled circles. The vertical offset of the circles indicates the deviation from the predicted values for that atom. Filled circles indicate that there are chemical shifts for atoms with the same set of attributes in the BMRB. Open circles indicate atoms that have a prediction, but for which no exact matches of the attributes are available in the BMRB. Grey boxes represent atoms that are not present in a given base.

**Figure 7.**
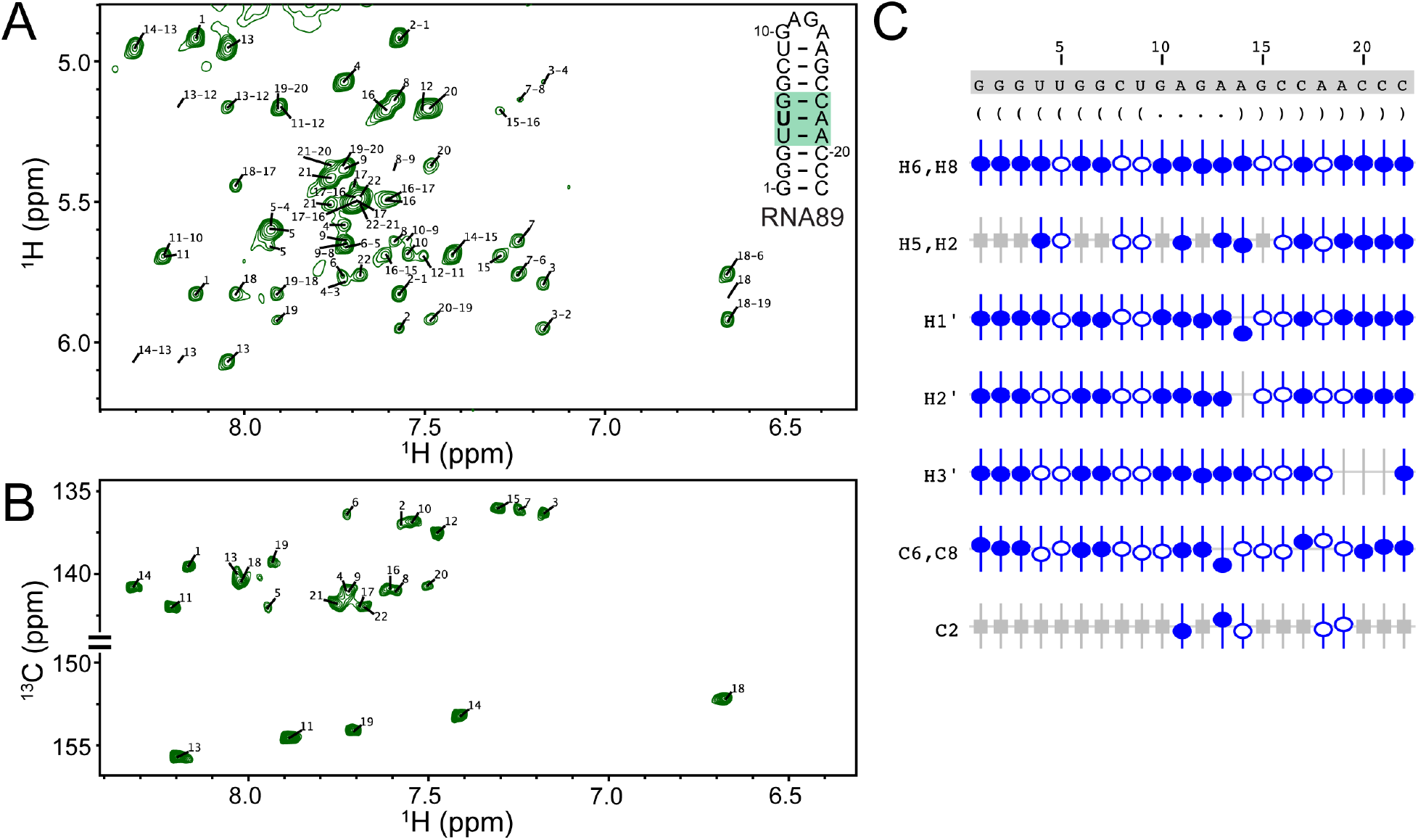
RNA89 chemical shift assignments. A) ^1^H-^1^H NOESY, B) ^1^H-^13^C HMQC, C) Summary of the sequence, secondary structure, and assignment validation for RNA89. The secondary structure is shown in Vienna format. NMRViewJ assignment summary to validate proton (H6/H8, H5/H2, H1′, H2′, H3′) and carbon (C6/C8, C2) assignments (bottom). Assigned chemical shifts for specific atoms are indicated with open and filled circles. The vertical offset of the circles indicates the deviation from the predicted values for that atom. Filled circles indicate that there are chemical shifts for atoms with the same set of attributes in the BMRB. Open circles indicate atoms that have a prediction, but for which no exact matches of the attributes are available in the BMRB. Grey boxes represent atoms that are not present in a given base.

Assigned chemical shifts along with raw NMR data have been deposited in the BMRB. RNA5: 50933, RNA7: 50932, RNA8: 50931, RNA21: 50930, RNA23: 50929, RNA24: 50928, RNA73: 50927, RNA74: 50926, RNA75: 50925, RNA89: 50924, RNA90: 50923, RNA91: 50922. The new data has also been used to update the training of chemical shift predictions in NMRFx Analyst (Marchant et al., 2019). The more complete database will immediately allow for better quality predictions in the use of the molecular network assignment tool integrated in NMRFx Analyst (Marchant et al., 2019). Similarly, these additional data will aid in machine learning approaches that predict RNA secondary structure (Zhang and Frank, 2020) and other RNA structural features from the assigned chemical shift data (Zhang et al., 2021).

## Acknowledgements

This work was supported by National Science Foundation grant MCB-1942398 (to S.C.K.) and by National Institute of General Medical Sciences of the National Institutes of Health grant U54 GM 103297 (to B.A.J.). Research reported in this publication was supported by the University of Michigan BioNMR Core Facility (U-M BioNMR). U-M BioNMR Core is grateful for support from U-M including the College of Literature, Sciences and Arts, Life Sciences Institute, College of Pharmacy and the Medical School along with the U-M Biosciences Initiative.

