## Supporting Information for "NMR chemical shift assignments of RNA oligonucleotides to expand the RNA chemical shift database"

[5] Current Address: Slovenian NMR Centre, National Institute of Chemistry, Hajdrihova 19, SI-1000 Ljubljana, Slovenia

**Table S1.** Biological RNAs with base pairing represented by the 12 RNAs investigated in this study.

| RNA# | Nucleotide numbers 5′→3′:3′→5′ | Biological RNA | PDB | Nucleotide numbers 5′→3′:3′→5′ |
|---|---|---|---|---|
| 5 | 1-6:17-22 | Ribosome | 1VW7 | 2122-2128:2326-2331 |
| | 1-5:18-22 | SAM-III riboswitch | 3E5C | 2-6:48-52 |
| 7 | 1-5:18-22 | HIV dimer initiation complex | 1BAU | 1-5:19-23 |
| | 2-7:16-21 | Ribosome | 5ANC | 2418-2423:2446-2452 |
| 8 | 3-7:16-20 | Ribosome | 1FFK | 906-910:1295-1299 |
| | 4-8:15-19 | HCV IRES | 1KH6 | 24-28:33-37 |
| 21 | 4-8:15-19 | Ribosome | 1FFK | 132-136:142-146 |
| | 5-9:14-18 | HIV (MAL) dimer initiation complex | 1K9W | 2-6:18-22 |
| 23 | 3-8:15-20 | Ribosome | 1VW7 | 2151-2156:2180-2186 |
| | 4-8:15-19 | Signal recognition particle RNA | 2XXA | 3-7:96-100 |
| 24 | 2-6:17-21 | tRNA Gly acceptor stem | 2VUQ | 1-5:68-72 |
| | 3-6:17-20 | Ribosome | 1VW7 | 2151-2154:2182-2185 |
| 73 | 2-8:15-21 | Pre-5S rRNA | 2I91 | (C)1-7:(D)2-8 |
| | 5-9:15-18 | Smaug response element RNA | 2ES5 | 5-9:15-19 |
| 74 | 2-7:16-21 | *E. coli* copy control RNA duplex | 1QC0 | (C)109-114:(D)125-130 |
| | 3-7:16-20 | 2'-Deoxyguanosine riboswitch | 3SKI | (A)35-39:47-51 |
| | 3-7:16-20 | Ribosome | 4V8P | 2649-2653:2695-2699 |
| | 3-7:16-20 | Ribosome | 1NJM | 2029-2033:2597-2601 |
| | 2-7:16-21 | *E. coli* copy control RNA duplex | 1QC0 | (C)109-114:(D)125-130 |
| 75 | 2-7:16-21 | Mitochondrial ribosome | 5AJ3 | 409-414:445-450 |
| | 3-6:17-20 | Ribonuclease P RNA | 1NBS | 92-95:108-111 |
| | 3-6:17-20 | Ribosome | 4L47 | 998-1001:1039-1042 |
| | 3-6:17-20 | MLV encapsidation signal | 16UP | 283-286:295-298 |
| 89 | 1-7:16-22 | Hairpin ribozyme | 1M5K | 29-35:46-52 |
| | 3-7:16-20 | *E. coli* copy control RNA duplex | 1QC0 | 106-110:129-133 |
| | 3-7:16-20 | Ribosome | 1FFK | 1734-1738:2041-2045 |
| | 3-6:17-20 | Ribosome | 3J3E | 3-6:110-113 |
| 90 | 2-7:16-21 | Selenocysteine insertion sequence RNA | 2RLU | 1-6:14-19 |
| | 3-8:15-20 | Cole1 inverted loop | 1BJ2 | 1-6:16-21 |
| | 3-8:15-20 | Selenocysteine insertion sequence RNA | 2PLY | 16-21:27-32 |
| | 3-7:16-20 | HIV Lai kissing complex | 2F4X | 2-6:20-24 |
| | 3-7:16-20 | tRNA | 1ZL3 | 402-406:418-422 |
| | 3-7:16-20 | Ribosome | 3J3E | 1428-1432:1503-1507 |
| 91 | 2-7:16-21 | PreQ1 riboswitch | 2MIY | -1-5:22-27 |
| | 3-7:16-20 | Ribosome | 3ZEX | 729-733:1587-1591 |
| | 3-7:16-20 | tRNA His | 3WC1 | 48-52:60-64 |
| | 3-7:16-20 | Hepatitis B virus encapsidation signal | 2IXY | 6-10:17-21 |
| | 3-7:16-20 | Ribosome | 4V7R | 2365-2369:2378-2382 |

**Table S2.** Summary of 5-base pair RNA sequences and their reported chemical shifts.

| | Sequence | Number of measured chemical shifts | | | | | | | | Complete chemical shift data? |
|---|---|---|---|---|---|---|---|---|---|---|
| | | H8 | C8 | H2 | C2 | H5 | C5 | H6 | C6 | |
| RNA1 | RY.AU.AU.AU.RY | 4 | 4 | 4 | 4 | 0 | 0 | 0 | 0 | TRUE |
| RNA2 | RY.AU.AU.AU.YR | 5 | 3 | 5 | 3 | 0 | 0 | 0 | 0 | TRUE |
| RNA3 | YR.AU.AU.AU.RY | 0 | 0 | 0 | 0 | 0 | 0 | 0 | 0 | FALSE |
| RNA4 | YR.AU.AU.AU.YR | 3 | 3 | 3 | 3 | 0 | 0 | 0 | 0 | TRUE |
| RNA5 | RY.AU.AU.UA.RY | 1 | 0 | 1 | 0 | 0 | 0 | 0 | 0 | FALSE |
| RNA6 | RY.AU.AU.UA.YR | 4 | 2 | 5 | 3 | 0 | 0 | 0 | 0 | TRUE |
| RNA7 | YR.AU.AU.UA.RY | 0 | 0 | 0 | 0 | 0 | 0 | 0 | 0 | FALSE |
| RNA8 | YR.AU.AU.UA.YR | 0 | 0 | 0 | 0 | 0 | 0 | 0 | 0 | FALSE |
| RNA9 | RY.AU.AU.GC.RY | 6 | 3 | 7 | 3 | 0 | 0 | 0 | 0 | TRUE |
| RNA10 | RY.AU.AU.GC.YR | 4 | 3 | 4 | 3 | 0 | 0 | 0 | 0 | TRUE |
| RNA11 | YR.AU.AU.GC.RY | 1 | 0 | 2 | 1 | 0 | 0 | 0 | 0 | FALSE |
| RNA12 | YR.AU.AU.GC.YR | 1 | 1 | 1 | 1 | 0 | 0 | 0 | 0 | TRUE |
| RNA13 | RY.AU.AU.CG.RY | 5 | 4 | 5 | 4 | 0 | 0 | 0 | 0 | TRUE |
| RNA14 | RY.AU.AU.CG.YR | 5 | 4 | 5 | 4 | 0 | 0 | 0 | 0 | TRUE |
| RNA15 | YR.AU.AU.CG.RY | 2 | 1 | 2 | 1 | 0 | 0 | 0 | 0 | TRUE |
| RNA16 | YR.AU.AU.CG.YR | 1 | 0 | 1 | 0 | 0 | 0 | 0 | 0 | FALSE |
| RNA17 | RY.UA.AU.AU.RY | 1 | 0 | 1 | 0 | 0 | 0 | 0 | 0 | FALSE |
| RNA18 | RY.UA.AU.AU.YR | 0 | 0 | 0 | 0 | 0 | 0 | 0 | 0 | FALSE |
| RNA19 | YR.UA.AU.AU.RY | 1 | 1 | 1 | 1 | 0 | 0 | 0 | 0 | TRUE |
| RNA20 | YR.UA.AU.AU.YR | 2 | 1 | 2 | 1 | 0 | 0 | 0 | 0 | TRUE |
| RNA21 | RY.UA.AU.UA.RY | 9 | 5 | 9 | 8 | 0 | 0 | 0 | 0 | TRUE |
| RNA22 | RY.UA.AU.UA.YR | 6 | 4 | 6 | 4 | 0 | 0 | 0 | 0 | TRUE |
| RNA23 | YR.UA.AU.UA.RY | 3 | 2 | 3 | 3 | 0 | 0 | 0 | 0 | TRUE |
| RNA24 | YR.UA.AU.UA.YR | 1 | 0 | 1 | 0 | 0 | 0 | 0 | 0 | FALSE |
| RNA25 | RY.UA.AU.GC.RY | 1 | 0 | 1 | 0 | 0 | 0 | 0 | 0 | FALSE |
| RNA26 | RY.UA.AU.GC.YR | 1 | 0 | 1 | 0 | 0 | 0 | 0 | 0 | FALSE |
| RNA27 | YR.UA.AU.GC.RY | 0 | 0 | 0 | 0 | 0 | 0 | 0 | 0 | FALSE |
| RNA28 | YR.UA.AU.GC.YR | 7 | 2 | 7 | 2 | 0 | 0 | 0 | 0 | TRUE |
| RNA29 | RY.UA.AU.CG.RY | 0 | 0 | 0 | 0 | 0 | 0 | 0 | 0 | FALSE |
| RNA30 | RY.UA.AU.CG.YR | 6 | 5 | 6 | 6 | 0 | 0 | 0 | 0 | TRUE |
| RNA31 | YR.UA.AU.CG.RY | 1 | 1 | 1 | 1 | 0 | 0 | 0 | 0 | TRUE |
| RNA32 | YR.UA.AU.CG.YR | 1 | 1 | 1 | 1 | 0 | 0 | 0 | 0 | TRUE |
| RNA33 | RY.GC.AU.AU.RY | 9 | 6 | 10 | 6 | 0 | 0 | 0 | 0 | TRUE |
| RNA34 | RY.GC.AU.AU.YR | 7 | 3 | 7 | 4 | 0 | 0 | 0 | 0 | TRUE |
| RNA35 | YR.GC.AU.AU.RY | 2 | 0 | 2 | 0 | 0 | 0 | 0 | 0 | FALSE |
| RNA36 | YR.GC.AU.AU.YR | 0 | 0 | 0 | 0 | 0 | 0 | 0 | 0 | FALSE |
| RNA37 | RY.GC.AU.UA.RY | 2 | 2 | 2 | 2 | 0 | 0 | 0 | 0 | TRUE |
| RNA38 | RY.GC.AU.UA.YR | 8 | 4 | 9 | 4 | 0 | 0 | 0 | 0 | TRUE |
| RNA39 | YR.GC.AU.UA.RY | 3 | 2 | 3 | 2 | 0 | 0 | 0 | 0 | TRUE |
| RNA40 | YR.GC.AU.UA.YR | 5 | 3 | 6 | 3 | 0 | 0 | 0 | 0 | TRUE |
| RNA41 | RY.GC.AU.GC.RY | 8 | 4 | 9 | 4 | 0 | 0 | 0 | 0 | TRUE |

| | | | | | | | | | | |
|---|---|---|---|---|---|---|---|---|---|---|
| RNA42 | RY.GC.AU.GC.YR | 13 | 10 | 13 | 12 | 0 | 0 | 0 | 0 | TRUE |
| RNA43 | YR.GC.AU.GC.RY | 5 | 1 | 5 | 1 | 0 | 0 | 0 | 0 | TRUE |
| RNA44 | YR.GC.AU.GC.YR | 2 | 1 | 2 | 1 | 0 | 0 | 0 | 0 | TRUE |
| RNA45 | RY.GC.AU.CG.RY | 8 | 7 | 8 | 6 | 0 | 0 | 0 | 0 | TRUE |
| RNA46 | RY.GC.AU.CG.YR | 13 | 6 | 12 | 5 | 0 | 0 | 0 | 0 | TRUE |
| RNA47 | YR.GC.AU.CG.RY | 0 | 0 | 0 | 0 | 0 | 0 | 0 | 0 | FALSE |
| RNA48 | YR.GC.AU.CG.YR | 3 | 2 | 3 | 2 | 0 | 0 | 0 | 0 | TRUE |
| RNA49 | RY.CG.AU.AU.RY | 5 | 2 | 5 | 2 | 0 | 0 | 0 | 0 | TRUE |
| RNA50 | RY.CG.AU.AU.YR | 2 | 2 | 3 | 2 | 0 | 0 | 0 | 0 | TRUE |
| RNA51 | YR.CG.AU.AU.RY | 2 | 2 | 1 | 1 | 0 | 0 | 0 | 0 | TRUE |
| RNA52 | YR.CG.AU.AU.YR | 0 | 0 | 0 | 0 | 0 | 0 | 0 | 0 | FALSE |
| RNA53 | RY.CG.AU.UA.RY | 7 | 5 | 7 | 6 | 0 | 0 | 0 | 0 | TRUE |
| RNA54 | RY.CG.AU.UA.YR | 0 | 0 | 0 | 0 | 0 | 0 | 0 | 0 | FALSE |
| RNA55 | YR.CG.AU.UA.RY | 0 | 0 | 0 | 0 | 0 | 0 | 0 | 0 | FALSE |
| RNA56 | YR.CG.AU.UA.YR | 4 | 4 | 4 | 4 | 0 | 0 | 0 | 0 | TRUE |
| RNA57 | RY.CG.AU.GC.RY | 9 | 3 | 9 | 3 | 0 | 0 | 0 | 0 | TRUE |
| RNA58 | RY.CG.AU.GC.YR | 8 | 8 | 7 | 7 | 0 | 0 | 0 | 0 | TRUE |
| RNA59 | YR.CG.AU.GC.RY | 7 | 3 | 7 | 1 | 0 | 0 | 0 | 0 | TRUE |
| RNA60 | YR.CG.AU.GC.YR | 5 | 3 | 5 | 2 | 0 | 0 | 0 | 0 | TRUE |
| RNA61 | RY.CG.AU.CG.RY | 6 | 5 | 6 | 5 | 0 | 0 | 0 | 0 | TRUE |
| RNA62 | RY.CG.AU.CG.YR | 13 | 10 | 13 | 10 | 0 | 0 | 0 | 0 | TRUE |
| RNA63 | YR.CG.AU.CG.RY | 0 | 0 | 0 | 0 | 0 | 0 | 0 | 0 | FALSE |
| RNA64 | YR.CG.AU.CG.YR | 4 | 1 | 4 | 1 | 0 | 0 | 0 | 0 | TRUE |
| RNA65 | RY.AU.UA.AU.RY | 0 | 0 | 0 | 0 | 1 | 0 | 1 | 0 | FALSE |
| RNA66 | RY.AU.UA.AU.YR | 0 | 0 | 0 | 0 | 3 | 2 | 5 | 2 | TRUE |
| RNA67 | YR.AU.UA.AU.RY | 0 | 0 | 0 | 0 | 1 | 1 | 2 | 0 | FALSE |
| RNA68 | YR.AU.UA.AU.YR | 0 | 0 | 0 | 0 | 7 | 4 | 8 | 3 | TRUE |
| RNA69 | RY.AU.UA.UA.RY | 0 | 0 | 0 | 0 | 0 | 0 | 0 | 0 | FALSE |
| RNA70 | RY.AU.UA.UA.YR | 0 | 0 | 0 | 0 | 2 | 2 | 4 | 1 | TRUE |
| RNA71 | YR.AU.UA.UA.RY | 0 | 0 | 0 | 0 | 0 | 0 | 0 | 0 | FALSE |
| RNA72 | YR.AU.UA.UA.YR | 0 | 0 | 0 | 0 | 0 | 0 | 1 | 0 | FALSE |
| RNA73 | RY.AU.UA.GC.RY | 0 | 0 | 0 | 0 | 3 | 3 | 3 | 3 | TRUE |
| RNA74 | RY.AU.UA.GC.YR | 0 | 0 | 0 | 0 | 0 | 0 | 0 | 0 | FALSE |
| RNA75 | YR.AU.UA.GC.RY | 0 | 0 | 0 | 0 | 0 | 0 | 0 | 0 | FALSE |
| RNA76 | YR.AU.UA.GC.YR | 0 | 0 | 0 | 0 | 7 | 7 | 7 | 6 | TRUE |
| RNA77 | RY.AU.UA.CG.RY | 0 | 0 | 0 | 0 | 8 | 4 | 8 | 7 | TRUE |
| RNA78 | RY.AU.UA.CG.YR | 0 | 0 | 0 | 0 | 11 | 9 | 12 | 8 | TRUE |
| RNA79 | YR.AU.UA.CG.RY | 0 | 0 | 0 | 0 | 4 | 3 | 4 | 3 | TRUE |
| RNA80 | YR.AU.UA.CG.YR | 0 | 0 | 0 | 0 | 2 | 2 | 2 | 2 | TRUE |
| RNA81 | RY.UA.UA.AU.RY | 0 | 0 | 0 | 0 | 2 | 1 | 2 | 1 | TRUE |
| RNA82 | RY.UA.UA.AU.YR | 0 | 0 | 0 | 0 | 0 | 0 | 0 | 0 | FALSE |
| RNA83 | YR.UA.UA.AU.RY | 0 | 0 | 0 | 0 | 1 | 1 | 1 | 1 | TRUE |
| RNA84 | YR.UA.UA.AU.YR | 0 | 0 | 0 | 0 | 1 | 0 | 1 | 0 | FALSE |
| RNA85 | RY.UA.UA.UA.RY | 0 | 0 | 0 | 0 | 2 | 2 | 2 | 2 | TRUE |

| | | | | | | | | | | |
|---|---|---|---|---|---|---|---|---|---|---|
| RNA86 | RY.UA.UA.UA.YR | 0 | 0 | 0 | 0 | 5 | 3 | 5 | 3 | TRUE |
| RNA87 | YR.UA.UA.UA.RY | 0 | 0 | 0 | 0 | 0 | 0 | 0 | 0 | FALSE |
| RNA88 | YR.UA.UA.UA.YR | 0 | 0 | 0 | 0 | 5 | 3 | 5 | 4 | TRUE |
| RNA89 | RY.UA.UA.GC.RY | 0 | 0 | 0 | 0 | 0 | 0 | 0 | 0 | FALSE |
| RNA90 | RY.UA.UA.GC.YR | 0 | 0 | 0 | 0 | 1 | 0 | 1 | 0 | FALSE |
| RNA91 | YR.UA.UA.GC.RY | 0 | 0 | 0 | 0 | 2 | 0 | 3 | 1 | FALSE |
| RNA92 | YR.UA.UA.GC.YR | 0 | 0 | 0 | 0 | 5 | 2 | 5 | 2 | TRUE |
| RNA93 | RY.UA.UA.CG.RY | 0 | 0 | 0 | 0 | 0 | 0 | 0 | 0 | FALSE |
| RNA94 | RY.UA.UA.CG.YR | 0 | 0 | 0 | 0 | 5 | 4 | 6 | 3 | TRUE |
| RNA95 | YR.UA.UA.CG.RY | 0 | 0 | 0 | 0 | 4 | 0 | 4 | 1 | FALSE |
| RNA96 | YR.UA.UA.CG.YR | 0 | 0 | 0 | 0 | 10 | 7 | 10 | 6 | TRUE |
| RNA97 | RY.GC.UA.AU.RY | 0 | 0 | 0 | 0 | 1 | 1 | 1 | 1 | TRUE |
| RNA98 | RY.GC.UA.AU.YR | 0 | 0 | 0 | 0 | 6 | 5 | 6 | 4 | TRUE |
| RNA99 | YR.GC.UA.AU.RY | 0 | 0 | 0 | 0 | 1 | 0 | 1 | 0 | FALSE |
| RNA100 | YR.GC.UA.AU.YR | 0 | 0 | 0 | 0 | 1 | 1 | 1 | 1 | TRUE |
| RNA101 | RY.GC.UA.UA.RY | 0 | 0 | 0 | 0 | 1 | 0 | 1 | 0 | FALSE |
| RNA102 | RY.GC.UA.UA.YR | 0 | 0 | 0 | 0 | 4 | 3 | 4 | 3 | TRUE |
| RNA103 | YR.GC.UA.UA.RY | 0 | 0 | 0 | 0 | 2 | 1 | 2 | 1 | TRUE |
| RNA104 | YR.GC.UA.UA.YR | 0 | 0 | 0 | 0 | 4 | 4 | 5 | 4 | TRUE |
| RNA105 | RY.GC.UA.GC.RY | 0 | 0 | 0 | 0 | 5 | 1 | 5 | 1 | TRUE |
| RNA106 | RY.GC.UA.GC.YR | 0 | 0 | 0 | 0 | 12 | 9 | 12 | 9 | TRUE |
| RNA107 | YR.GC.UA.GC.RY | 0 | 0 | 0 | 0 | 2 | 2 | 2 | 2 | TRUE |
| RNA108 | YR.GC.UA.GC.YR | 0 | 0 | 0 | 0 | 3 | 3 | 3 | 3 | TRUE |
| RNA109 | RY.GC.UA.CG.RY | 0 | 0 | 0 | 0 | 2 | 1 | 2 | 1 | TRUE |
| RNA110 | RY.GC.UA.CG.YR | 0 | 0 | 0 | 0 | 15 | 8 | 16 | 9 | TRUE |
| RNA111 | YR.GC.UA.CG.RY | 0 | 0 | 0 | 0 | 0 | 0 | 0 | 0 | FALSE |
| RNA112 | YR.GC.UA.CG.YR | 0 | 0 | 0 | 0 | 6 | 6 | 6 | 6 | TRUE |
| RNA113 | RY.CG.UA.AU.RY | 0 | 0 | 0 | 0 | 7 | 2 | 7 | 2 | TRUE |
| RNA114 | RY.CG.UA.AU.YR | 0 | 0 | 0 | 0 | 1 | 1 | 1 | 1 | TRUE |
| RNA115 | YR.CG.UA.AU.RY | 0 | 0 | 0 | 0 | 0 | 0 | 0 | 0 | FALSE |
| RNA116 | YR.CG.UA.AU.YR | 0 | 0 | 0 | 0 | 1 | 1 | 1 | 1 | TRUE |
| RNA117 | RY.CG.UA.UA.RY | 0 | 0 | 0 | 0 | 1 | 0 | 1 | 1 | FALSE |
| RNA118 | RY.CG.UA.UA.YR | 0 | 0 | 0 | 0 | 4 | 3 | 4 | 2 | TRUE |
| RNA119 | YR.CG.UA.UA.RY | 0 | 0 | 0 | 0 | 1 | 0 | 3 | 0 | FALSE |
| RNA120 | YR.CG.UA.UA.YR | 0 | 0 | 0 | 0 | 6 | 4 | 6 | 2 | TRUE |
| RNA121 | RY.CG.UA.GC.RY | 0 | 0 | 0 | 0 | 4 | 2 | 4 | 3 | TRUE |
| RNA122 | RY.CG.UA.GC.YR | 0 | 0 | 0 | 0 | 5 | 5 | 5 | 5 | TRUE |
| RNA123 | YR.CG.UA.GC.RY | 0 | 0 | 0 | 0 | 9 | 6 | 9 | 5 | TRUE |
| RNA124 | YR.CG.UA.GC.YR | 0 | 0 | 0 | 0 | 11 | 5 | 11 | 5 | TRUE |
| RNA125 | RY.CG.UA.CG.RY | 0 | 0 | 0 | 0 | 2 | 1 | 2 | 1 | TRUE |
| RNA126 | RY.CG.UA.CG.YR | 0 | 0 | 0 | 0 | 16 | 12 | 16 | 12 | TRUE |
| RNA127 | YR.CG.UA.CG.RY | 0 | 0 | 0 | 0 | 2 | 0 | 2 | 0 | FALSE |
| RNA128 | YR.CG.UA.CG.YR | 0 | 0 | 0 | 0 | 15 | 7 | 15 | 8 | TRUE |
| RNA129 | RY.AU.GC.AU.RY | 2 | 1 | 0 | 0 | 0 | 0 | 0 | 0 | TRUE |

| | | | | | | | | | | |
|---|---|---|---|---|---|---|---|---|---|---|
| RNA130 | RY.AU.GC.AU.YR | 3 | 2 | 0 | 0 | 0 | 0 | 0 | 0 | TRUE |
| RNA131 | YR.AU.GC.AU.RY | 3 | 1 | 0 | 0 | 0 | 0 | 0 | 0 | TRUE |
| RNA132 | YR.AU.GC.AU.YR | 2 | 1 | 0 | 0 | 0 | 0 | 0 | 0 | TRUE |
| RNA133 | RY.AU.GC.UA.RY | 9 | 9 | 0 | 0 | 0 | 0 | 0 | 0 | TRUE |
| RNA134 | RY.AU.GC.UA.YR | 3 | 0 | 0 | 0 | 0 | 0 | 0 | 0 | FALSE |
| RNA135 | YR.AU.GC.UA.RY | 3 | 1 | 0 | 0 | 0 | 0 | 0 | 0 | TRUE |
| RNA136 | YR.AU.GC.UA.YR | 4 | 2 | 0 | 0 | 0 | 0 | 0 | 0 | TRUE |
| RNA137 | RY.AU.GC.GC.RY | 7 | 2 | 0 | 0 | 0 | 0 | 0 | 0 | TRUE |
| RNA138 | RY.AU.GC.GC.YR | 9 | 6 | 0 | 0 | 0 | 0 | 0 | 0 | TRUE |
| RNA139 | YR.AU.GC.GC.RY | 1 | 0 | 0 | 0 | 0 | 0 | 0 | 0 | FALSE |
| RNA140 | YR.AU.GC.GC.YR | 2 | 2 | 0 | 0 | 0 | 0 | 0 | 0 | TRUE |
| RNA141 | RY.AU.GC.CG.RY | 6 | 4 | 0 | 0 | 0 | 0 | 0 | 0 | TRUE |
| RNA142 | RY.AU.GC.CG.YR | 8 | 7 | 0 | 0 | 0 | 0 | 0 | 0 | TRUE |
| RNA143 | YR.AU.GC.CG.RY | 0 | 0 | 0 | 0 | 0 | 0 | 0 | 0 | FALSE |
| RNA144 | YR.AU.GC.CG.YR | 9 | 6 | 0 | 0 | 0 | 0 | 0 | 0 | TRUE |
| RNA145 | RY.UA.GC.AU.RY | 2 | 0 | 0 | 0 | 0 | 0 | 0 | 0 | FALSE |
| RNA146 | RY.UA.GC.AU.YR | 1 | 1 | 0 | 0 | 0 | 0 | 0 | 0 | TRUE |
| RNA147 | YR.UA.GC.AU.RY | 2 | 2 | 0 | 0 | 0 | 0 | 0 | 0 | TRUE |
| RNA148 | YR.UA.GC.AU.YR | 3 | 0 | 0 | 0 | 0 | 0 | 0 | 0 | FALSE |
| RNA149 | RY.UA.GC.UA.RY | 2 | 1 | 0 | 0 | 0 | 0 | 0 | 0 | TRUE |
| RNA150 | RY.UA.GC.UA.YR | 6 | 4 | 0 | 0 | 0 | 0 | 0 | 0 | TRUE |
| RNA151 | YR.UA.GC.UA.RY | 0 | 0 | 0 | 0 | 0 | 0 | 0 | 0 | FALSE |
| RNA152 | YR.UA.GC.UA.YR | 2 | 2 | 0 | 0 | 0 | 0 | 0 | 0 | TRUE |
| RNA153 | RY.UA.GC.GC.RY | 3 | 2 | 0 | 0 | 0 | 0 | 0 | 0 | TRUE |
| RNA154 | RY.UA.GC.GC.YR | 6 | 5 | 0 | 0 | 0 | 0 | 0 | 0 | TRUE |
| RNA155 | YR.UA.GC.GC.RY | 3 | 1 | 0 | 0 | 0 | 0 | 0 | 0 | TRUE |
| RNA156 | YR.UA.GC.GC.YR | 7 | 6 | 0 | 0 | 0 | 0 | 0 | 0 | TRUE |
| RNA157 | RY.UA.GC.CG.RY | 2 | 0 | 0 | 0 | 0 | 0 | 0 | 0 | FALSE |
| RNA158 | RY.UA.GC.CG.YR | 6 | 5 | 0 | 0 | 0 | 0 | 0 | 0 | TRUE |
| RNA159 | YR.UA.GC.CG.RY | 1 | 0 | 0 | 0 | 0 | 0 | 0 | 0 | FALSE |
| RNA160 | YR.UA.GC.CG.YR | 18 | 8 | 0 | 0 | 0 | 0 | 0 | 0 | TRUE |
| RNA161 | RY.GC.GC.AU.RY | 11 | 7 | 0 | 0 | 0 | 0 | 0 | 0 | TRUE |
| RNA162 | RY.GC.GC.AU.YR | 10 | 6 | 0 | 0 | 0 | 0 | 0 | 0 | TRUE |
| RNA163 | YR.GC.GC.AU.RY | 4 | 2 | 0 | 0 | 0 | 0 | 0 | 0 | TRUE |
| RNA164 | YR.GC.GC.AU.YR | 1 | 0 | 0 | 0 | 0 | 0 | 0 | 0 | FALSE |
| RNA165 | RY.GC.GC.UA.RY | 7 | 3 | 0 | 0 | 0 | 0 | 0 | 0 | TRUE |
| RNA166 | RY.GC.GC.UA.YR | 6 | 2 | 0 | 0 | 0 | 0 | 0 | 0 | TRUE |
| RNA167 | YR.GC.GC.UA.RY | 3 | 3 | 0 | 0 | 0 | 0 | 0 | 0 | TRUE |
| RNA168 | YR.GC.GC.UA.YR | 2 | 3 | 0 | 0 | 0 | 0 | 0 | 0 | TRUE |
| RNA169 | RY.GC.GC.GC.RY | 8 | 7 | 0 | 0 | 0 | 0 | 0 | 0 | TRUE |
| RNA170 | RY.GC.GC.GC.YR | 3 | 0 | 0 | 0 | 0 | 0 | 0 | 0 | FALSE |
| RNA171 | YR.GC.GC.GC.RY | 6 | 2 | 0 | 0 | 0 | 0 | 0 | 0 | TRUE |
| RNA172 | YR.GC.GC.GC.YR | 1 | 0 | 0 | 0 | 0 | 0 | 0 | 0 | FALSE |
| RNA173 | RY.GC.GC.CG.RY | 5 | 3 | 0 | 0 | 0 | 0 | 0 | 0 | TRUE |

| | | | | | | | | | | |
|---|---|---|---|---|---|---|---|---|---|---|
| RNA174 | RY.GC.GC.CG.YR | 12 | 8 | 0 | 0 | 0 | 0 | 0 | 0 | TRUE |
| RNA175 | YR.GC.GC.CG.RY | 1 | 0 | 0 | 0 | 0 | 0 | 0 | 0 | FALSE |
| RNA176 | YR.GC.GC.CG.YR | 4 | 3 | 0 | 0 | 0 | 0 | 0 | 0 | TRUE |
| RNA177 | RY.CG.GC.AU.RY | 1 | 0 | 0 | 0 | 0 | 0 | 0 | 0 | FALSE |
| RNA178 | RY.CG.GC.AU.YR | 6 | 6 | 0 | 0 | 0 | 0 | 0 | 0 | TRUE |
| RNA179 | YR.CG.GC.AU.RY | 3 | 1 | 0 | 0 | 0 | 0 | 0 | 0 | TRUE |
| RNA180 | YR.CG.GC.AU.YR | 3 | 2 | 0 | 0 | 0 | 0 | 0 | 0 | TRUE |
| RNA181 | RY.CG.GC.UA.RY | 1 | 1 | 0 | 0 | 0 | 0 | 0 | 0 | TRUE |
| RNA182 | RY.CG.GC.UA.YR | 3 | 2 | 0 | 0 | 0 | 0 | 0 | 0 | TRUE |
| RNA183 | YR.CG.GC.UA.RY | 4 | 4 | 0 | 0 | 0 | 0 | 0 | 0 | TRUE |
| RNA184 | YR.CG.GC.UA.YR | 3 | 3 | 0 | 0 | 0 | 0 | 0 | 0 | TRUE |
| RNA185 | RY.CG.GC.GC.RY | 2 | 1 | 0 | 0 | 0 | 0 | 0 | 0 | TRUE |
| RNA186 | RY.CG.GC.GC.YR | 3 | 2 | 0 | 0 | 0 | 0 | 0 | 0 | TRUE |
| RNA187 | YR.CG.GC.GC.RY | 4 | 0 | 0 | 0 | 0 | 0 | 0 | 0 | FALSE |
| RNA188 | YR.CG.GC.GC.YR | 3 | 0 | 0 | 0 | 0 | 0 | 0 | 0 | FALSE |
| RNA189 | RY.CG.GC.CG.RY | 3 | 1 | 0 | 0 | 0 | 0 | 0 | 0 | TRUE |
| RNA190 | RY.CG.GC.CG.YR | 2 | 2 | 0 | 0 | 0 | 0 | 0 | 0 | TRUE |
| RNA191 | YR.CG.GC.CG.RY | 0 | 0 | 0 | 0 | 0 | 0 | 0 | 0 | FALSE |
| RNA192 | YR.CG.GC.CG.YR | 7 | 6 | 0 | 0 | 0 | 0 | 0 | 0 | TRUE |
| RNA193 | RY.AU.CG.AU.RY | 0 | 0 | 0 | 0 | 2 | 2 | 2 | 3 | TRUE |
| RNA194 | RY.AU.CG.AU.YR | 0 | 0 | 0 | 0 | 4 | 4 | 5 | 4 | TRUE |
| RNA195 | YR.AU.CG.AU.RY | 0 | 0 | 0 | 0 | 2 | 2 | 2 | 2 | TRUE |
| RNA196 | YR.AU.CG.AU.YR | 0 | 0 | 0 | 0 | 2 | 2 | 2 | 2 | TRUE |
| RNA197 | RY.AU.CG.UA.RY | 0 | 0 | 0 | 0 | 5 | 3 | 5 | 2 | TRUE |
| RNA198 | RY.AU.CG.UA.YR | 0 | 0 | 0 | 0 | 2 | 0 | 2 | 0 | FALSE |
| RNA199 | YR.AU.CG.UA.RY | 0 | 0 | 0 | 0 | 2 | 2 | 2 | 2 | TRUE |
| RNA200 | YR.AU.CG.UA.YR | 0 | 0 | 0 | 0 | 9 | 7 | 9 | 7 | TRUE |
| RNA201 | RY.AU.CG.GC.RY | 0 | 0 | 0 | 0 | 3 | 3 | 3 | 3 | TRUE |
| RNA202 | RY.AU.CG.GC.YR | 0 | 0 | 0 | 0 | 3 | 2 | 3 | 2 | TRUE |
| RNA203 | YR.AU.CG.GC.RY | 0 | 0 | 0 | 0 | 4 | 4 | 4 | 3 | TRUE |
| RNA204 | YR.AU.CG.GC.YR | 0 | 0 | 0 | 0 | 1 | 0 | 1 | 1 | FALSE |
| RNA205 | RY.AU.CG.CG.RY | 0 | 0 | 0 | 0 | 2 | 1 | 2 | 1 | TRUE |
| RNA206 | RY.AU.CG.CG.YR | 0 | 0 | 0 | 0 | 6 | 2 | 6 | 1 | TRUE |
| RNA207 | YR.AU.CG.CG.RY | 0 | 0 | 0 | 0 | 3 | 2 | 3 | 3 | TRUE |
| RNA208 | YR.AU.CG.CG.YR | 0 | 0 | 0 | 0 | 8 | 3 | 9 | 3 | TRUE |
| RNA209 | RY.UA.CG.AU.RY | 0 | 0 | 0 | 0 | 2 | 0 | 2 | 1 | FALSE |
| RNA210 | RY.UA.CG.AU.YR | 0 | 0 | 0 | 0 | 1 | 1 | 1 | 1 | TRUE |
| RNA211 | YR.UA.CG.AU.RY | 0 | 0 | 0 | 0 | 1 | 0 | 1 | 0 | FALSE |
| RNA212 | YR.UA.CG.AU.YR | 0 | 0 | 0 | 0 | 2 | 0 | 2 | 0 | FALSE |
| RNA213 | RY.UA.CG.UA.RY | 0 | 0 | 0 | 0 | 6 | 3 | 6 | 2 | TRUE |
| RNA214 | RY.UA.CG.UA.YR | 0 | 0 | 0 | 0 | 4 | 3 | 5 | 2 | TRUE |
| RNA215 | YR.UA.CG.UA.RY | 0 | 0 | 0 | 0 | 9 | 4 | 9 | 4 | TRUE |
| RNA216 | YR.UA.CG.UA.YR | 0 | 0 | 0 | 0 | 9 | 4 | 8 | 4 | TRUE |
| RNA217 | RY.UA.CG.GC.RY | 0 | 0 | 0 | 0 | 1 | 0 | 1 | 0 | FALSE |

| | | | | | | | | | | |
|---|---|---|---|---|---|---|---|---|---|---|
| RNA218 | RY.UA.CG.GC.YR | 0 | 0 | 0 | 0 | 8 | 5 | 8 | 5 | TRUE |
| RNA219 | YR.UA.CG.GC.RY | 0 | 0 | 0 | 0 | 2 | 1 | 2 | 1 | TRUE |
| RNA220 | YR.UA.CG.GC.YR | 0 | 0 | 0 | 0 | 1 | 0 | 1 | 0 | FALSE |
| RNA221 | RY.UA.CG.CG.RY | 0 | 0 | 0 | 0 | 0 | 0 | 0 | 0 | FALSE |
| RNA222 | RY.UA.CG.CG.YR | 0 | 0 | 0 | 0 | 12 | 6 | 12 | 6 | TRUE |
| RNA223 | YR.UA.CG.CG.RY | 0 | 0 | 0 | 0 | 3 | 1 | 3 | 1 | TRUE |
| RNA224 | YR.UA.CG.CG.YR | 0 | 0 | 0 | 0 | 12 | 7 | 12 | 6 | TRUE |
| RNA225 | RY.GC.CG.AU.RY | 0 | 0 | 0 | 0 | 17 | 6 | 18 | 6 | TRUE |
| RNA226 | RY.GC.CG.AU.YR | 0 | 0 | 0 | 0 | 6 | 4 | 6 | 5 | TRUE |
| RNA227 | YR.GC.CG.AU.RY | 0 | 0 | 0 | 0 | 2 | 0 | 2 | 0 | FALSE |
| RNA228 | YR.GC.CG.AU.YR | 0 | 0 | 0 | 0 | 1 | 0 | 1 | 0 | FALSE |
| RNA229 | RY.GC.CG.UA.RY | 0 | 0 | 0 | 0 | 6 | 3 | 6 | 3 | TRUE |
| RNA230 | RY.GC.CG.UA.YR | 0 | 0 | 0 | 0 | 6 | 6 | 6 | 6 | TRUE |
| RNA231 | YR.GC.CG.UA.RY | 0 | 0 | 0 | 0 | 2 | 2 | 2 | 2 | TRUE |
| RNA232 | YR.GC.CG.UA.YR | 0 | 0 | 0 | 0 | 6 | 3 | 6 | 3 | TRUE |
| RNA233 | RY.GC.CG.GC.RY | 0 | 0 | 0 | 0 | 3 | 1 | 3 | 1 | TRUE |
| RNA234 | RY.GC.CG.GC.YR | 0 | 0 | 0 | 0 | 2 | 1 | 2 | 1 | TRUE |
| RNA235 | YR.GC.CG.GC.RY | 0 | 0 | 0 | 0 | 1 | 1 | 1 | 1 | TRUE |
| RNA236 | YR.GC.CG.GC.YR | 0 | 0 | 0 | 0 | 2 | 0 | 2 | 1 | FALSE |
| RNA237 | RY.GC.CG.CG.RY | 0 | 0 | 0 | 0 | 6 | 4 | 6 | 4 | TRUE |
| RNA238 | RY.GC.CG.CG.YR | 0 | 0 | 0 | 0 | 12 | 8 | 11 | 7 | TRUE |
| RNA239 | YR.GC.CG.CG.RY | 0 | 0 | 0 | 0 | 1 | 0 | 1 | 0 | FALSE |
| RNA240 | YR.GC.CG.CG.YR | 0 | 0 | 0 | 0 | 6 | 3 | 7 | 3 | TRUE |
| RNA241 | RY.CG.CG.AU.RY | 0 | 0 | 0 | 0 | 6 | 2 | 6 | 5 | TRUE |
| RNA242 | RY.CG.CG.AU.YR | 0 | 0 | 0 | 0 | 3 | 4 | 5 | 3 | TRUE |
| RNA243 | YR.CG.CG.AU.RY | 0 | 0 | 0 | 0 | 3 | 1 | 3 | 2 | TRUE |
| RNA244 | YR.CG.CG.AU.YR | 0 | 0 | 0 | 0 | 2 | 2 | 2 | 1 | TRUE |
| RNA245 | RY.CG.CG.UA.RY | 0 | 0 | 0 | 0 | 3 | 2 | 3 | 1 | TRUE |
| RNA246 | RY.CG.CG.UA.YR | 0 | 0 | 0 | 0 | 8 | 4 | 9 | 3 | TRUE |
| RNA247 | YR.CG.CG.UA.RY | 0 | 0 | 0 | 0 | 0 | 0 | 0 | 0 | FALSE |
| RNA248 | YR.CG.CG.UA.YR | 0 | 0 | 0 | 0 | 7 | 3 | 7 | 3 | TRUE |
| RNA249 | RY.CG.CG.GC.RY | 0 | 0 | 0 | 0 | 3 | 2 | 3 | 2 | TRUE |
| RNA250 | RY.CG.CG.GC.YR | 0 | 0 | 0 | 0 | 3 | 2 | 3 | 2 | TRUE |
| RNA251 | YR.CG.CG.GC.RY | 0 | 0 | 0 | 0 | 4 | 1 | 4 | 0 | FALSE |
| RNA252 | YR.CG.CG.GC.YR | 0 | 0 | 0 | 0 | 2 | 2 | 2 | 2 | TRUE |
| RNA253 | RY.CG.CG.CG.RY | 0 | 0 | 0 | 0 | 7 | 0 | 7 | 0 | FALSE |
| RNA254 | RY.CG.CG.CG.YR | 0 | 0 | 0 | 0 | 3 | 1 | 3 | 1 | TRUE |
| RNA255 | YR.CG.CG.CG.RY | 0 | 0 | 0 | 0 | 5 | 0 | 5 | 2 | FALSE |
| RNA256 | YR.CG.CG.CG.YR | 0 | 0 | 0 | 0 | 11 | 5 | 11 | 7 | TRUE |

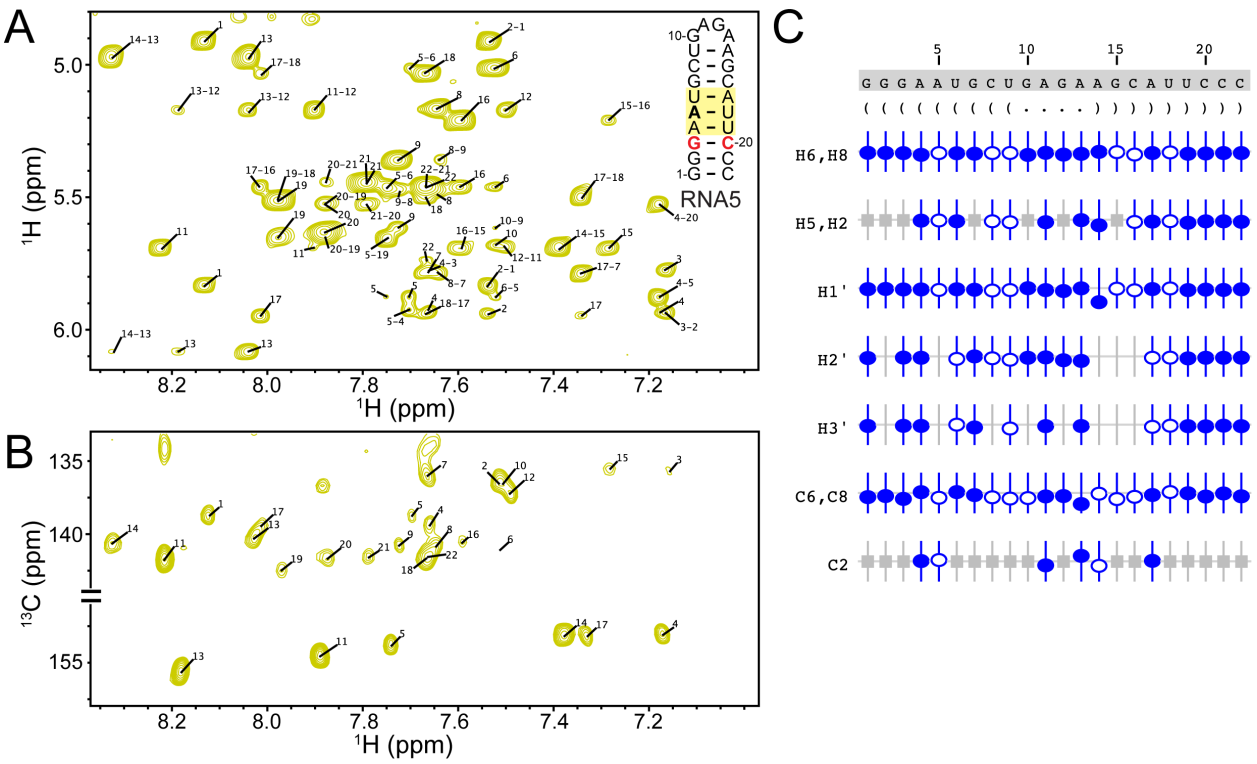


**Figure S1.** RNA5 chemical shift assignments. A) $^1$H-$^1$H NOESY, B) $^1$H-$^{13}$C HMQC, C) Summary of the sequence, secondary structure, and assignment validation for RNA5. The secondary structure is shown in Vienna format. NMRViewJ assignment summary to validate proton (H6/H8, H5/H2, H1′, H2′, H3′) and carbon (C6/C8, C2) assignments (bottom). Assigned chemical shifts for specific atoms are indicated with open and filled circles. The vertical offset of the circles indicates the deviation from the predicted values for that atom. Filled circles indicate that there are chemical shifts for atoms with the same set of attributes in the BMRB. Open circles indicate atoms that have a prediction, but for which no exact matches of the attributes are available in the BMRB. Grey boxes represent atoms that are not present in a given base.

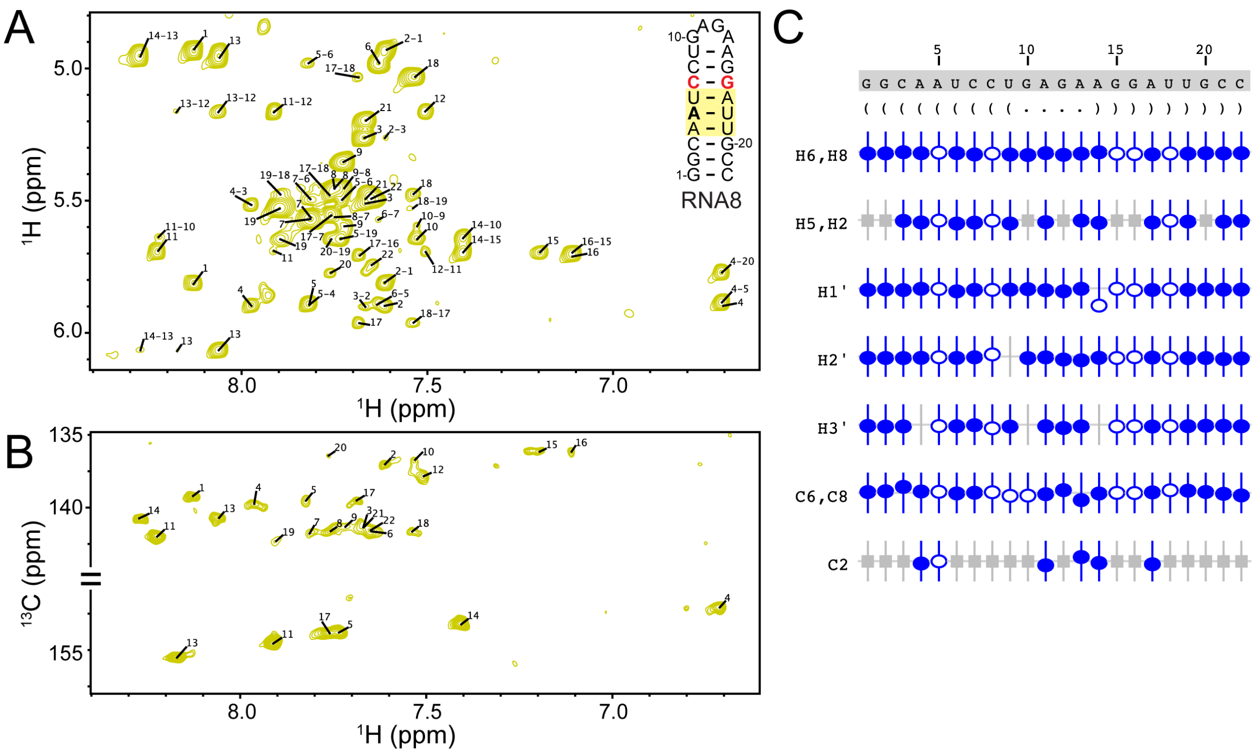


**Figure S2.** RNA8 chemical shift assignments. A) $^{1}$H-$^{1}$H NOESY, B) $^{1}$H-$^{13}$C HMQC, C) Summary of the sequence, secondary structure, and assignment validation for RNA8. The secondary structure is shown in Vienna format. NMRViewJ assignment summary to validate proton (H6/H8, H5/H2, H1′, H2′, H3′) and carbon (C6/C8, C2) assignments (bottom). Assigned chemical shifts for specific atoms are indicated with open and filled circles. The vertical offset of the circles indicates the deviation from the predicted values for that atom. Filled circles indicate that there are chemical shifts for atoms with the same set of attributes in the BMRB. Open circles indicate atoms that have a prediction, but for which no exact matches of the attributes are available in the BMRB. Grey boxes represent atoms that are not present in a given base.

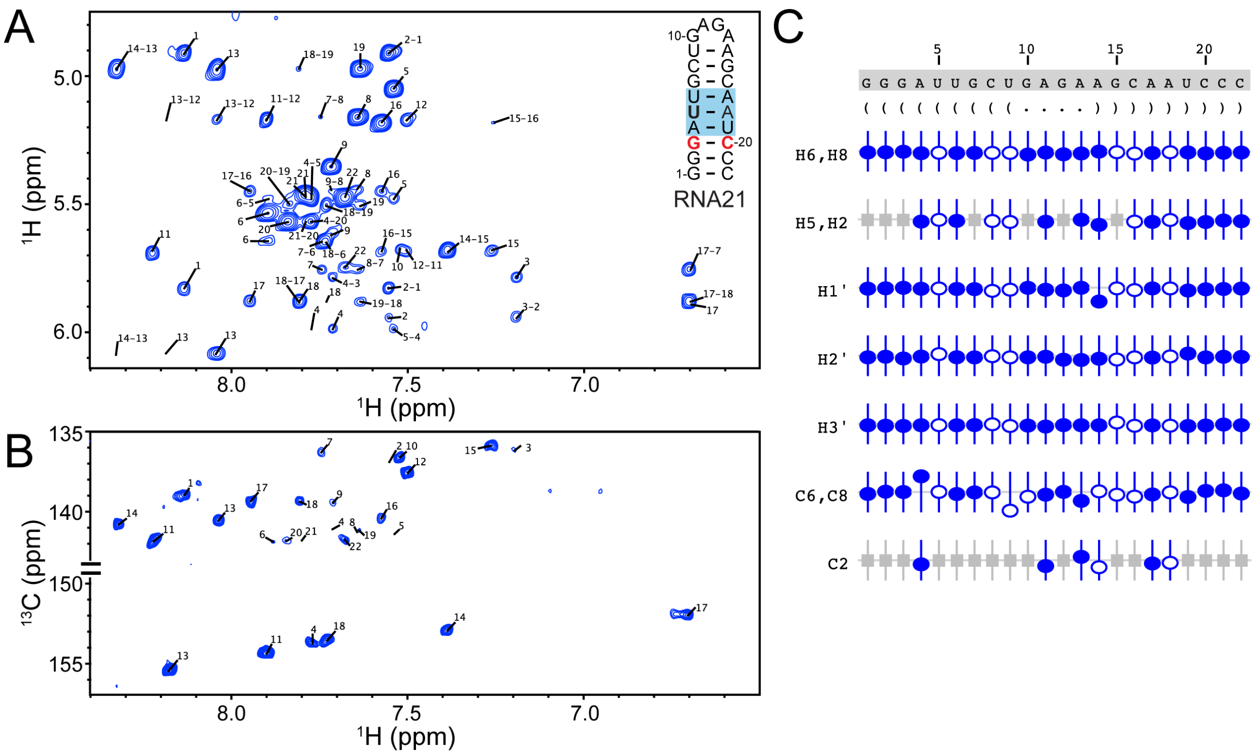


**Figure S3.** RNA21 chemical shift assignments. A) $^{1}$H-$^{1}$H NOESY, B) $^{1}$H-$^{13}$C HMQC, C) Summary of the sequence, secondary structure, and assignment validation for RNA21. The secondary structure is shown in Vienna format. NMRViewJ assignment summary to validate proton (H6/H8, H5/H2, H1′, H2′, H3′) and carbon (C6/C8, C2) assignments (bottom). Assigned chemical shifts for specific atoms are indicated with open and filled circles. The vertical offset of the circles indicates the deviation from the predicted values for that atom. Filled circles indicate that there are chemical shifts for atoms with the same set of attributes in the BMRB. Open circles indicate atoms that have a prediction, but for which no exact matches of the attributes are available in the BMRB. Grey boxes represent atoms that are not present in a given base.

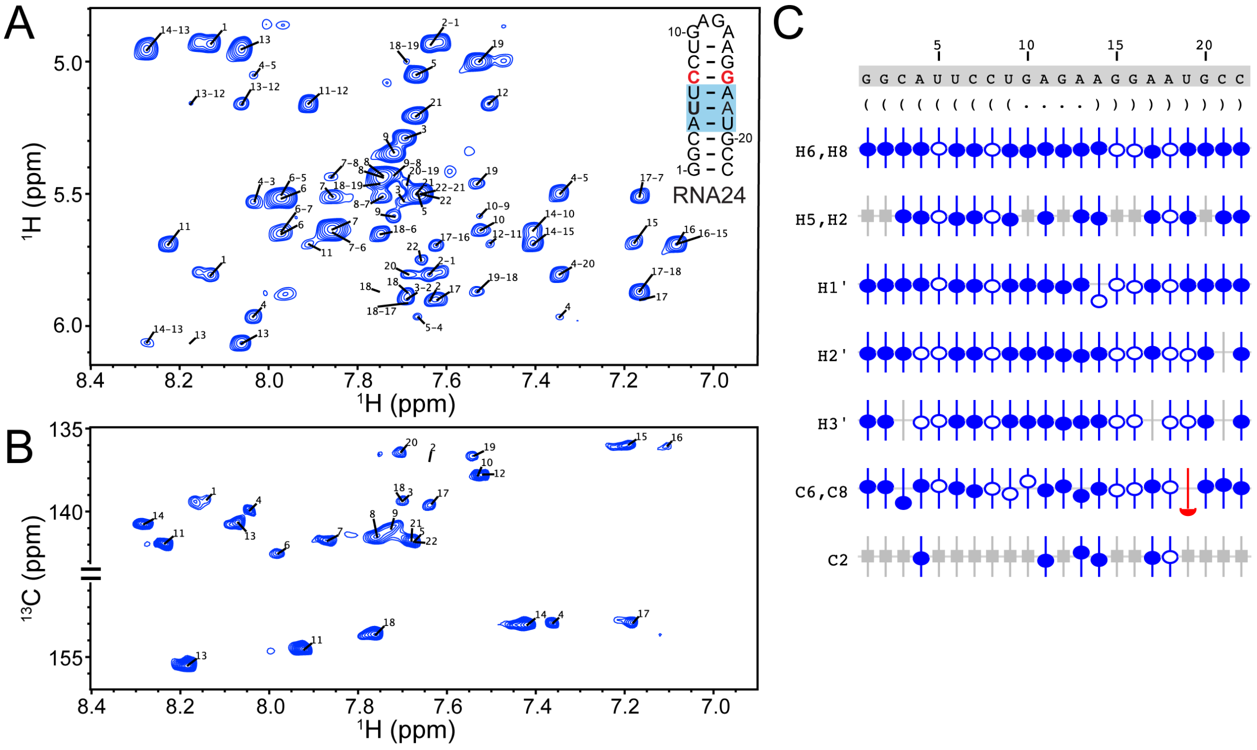


**Figure S4.** RNA24 chemical shift assignments. A) $^1$H-$^1$H NOESY, B) $^1$H-$^{13}$C HMQC, C) Summary of the sequence, secondary structure, and assignment validation for RNA24. The secondary structure is shown in Vienna format. NMRViewJ assignment summary to validate proton (H6/H8, H5/H2, H1′, H2′, H3′) and carbon (C6/C8, C2) assignments (bottom). Assigned chemical shifts for specific atoms are indicated with open and filled circles. The vertical offset of the circles indicates the deviation from the predicted values for that atom. Filled circles indicate that there are chemical shifts for atoms with the same set of attributes in the BMRB. Open circles indicate atoms that have a prediction, but for which no exact matches of the attributes are available in the BMRB. Red symbols indicate assignments that fall outside of the tolerance range. Grey boxes represent atoms that are not present in a given base.

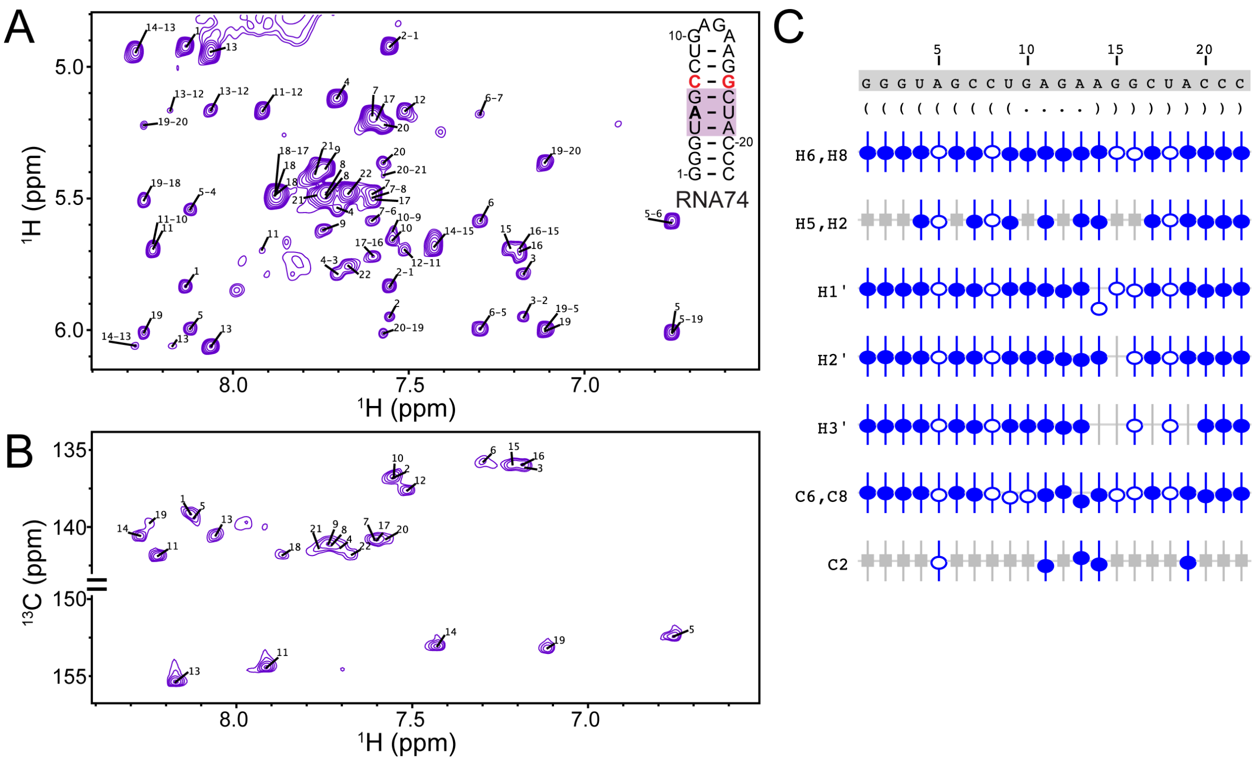


**Figure S5.** RNA74 chemical shift assignments. A) $^1$H-$^1$H NOESY, B) $^1$H-$^{13}$C HMQC, C) Summary of the sequence, secondary structure, and assignment validation for RNA74. The secondary structure is shown in Vienna format. NMRViewJ assignment summary to validate proton (H6/H8, H5/H2, H1′, H2′, H3′) and carbon (C6/C8, C2) assignments (bottom). Assignments are indicated with open and filled circles. Assigned chemical shifts for specific atoms are indicated with open and filled circles. The vertical offset of the circles indicates the deviation from the predicted values for that atom. Filled circles indicate that there are chemical shifts for atoms with the same set of attributes in the BMRB. Open circles indicate atoms that have a prediction, but for which no exact matches of the attributes are available in the BMRB. Grey boxes represent atoms that are not present in a given base.

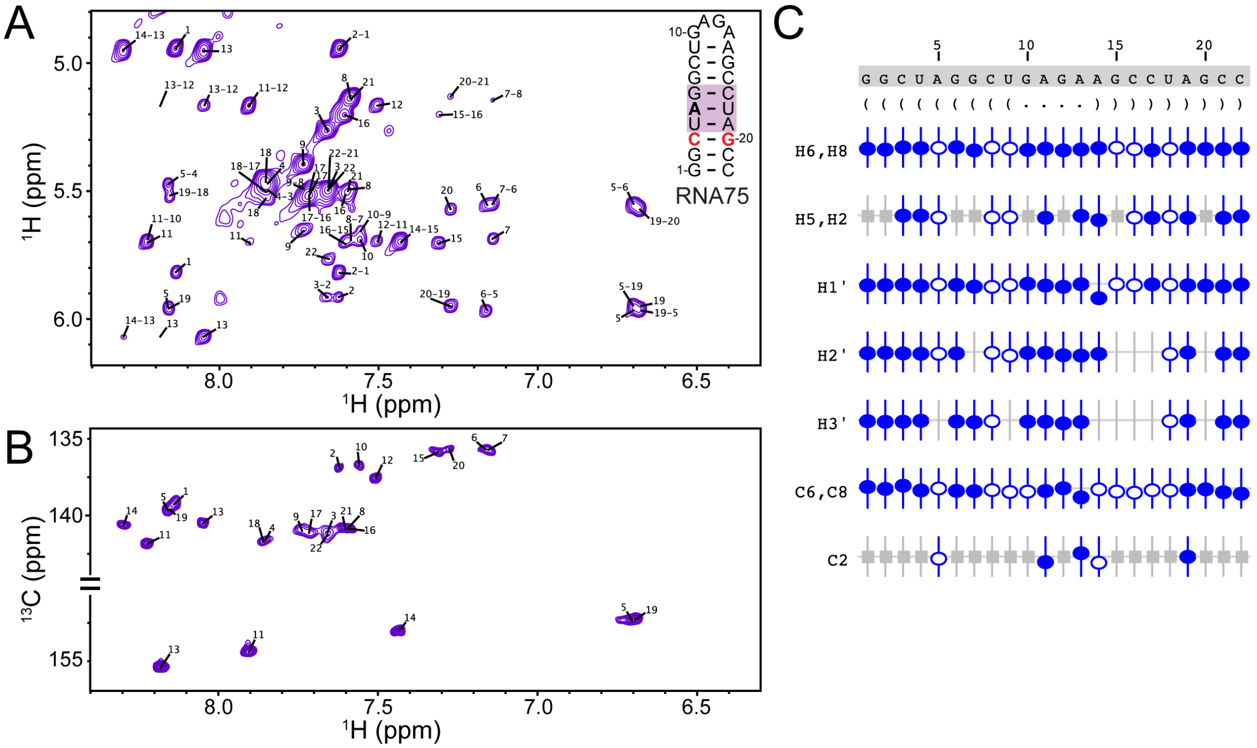


**Figure S6.** RNA75 chemical shift assignments. A) $^{1}$H-$^{1}$H NOESY, B) $^{1}$H-$^{13}$C HMQC, C) Summary of the sequence, secondary structure, and assignment validation for RNA75. The secondary structure is shown in Vienna format. NMRViewJ assignment summary to validate proton (H6/H8, H5/H2, H1′, H2′, H3′) and carbon (C6/C8, C2) assignments (bottom). Assignments are indicated with open and filled circles. Assigned chemical shifts for specific atoms are indicated with open and filled circles. The vertical offset of the circles indicates the deviation from the predicted values for that atom. Filled circles indicate that there are chemical shifts for atoms with the same set of attributes in the BMRB. Open circles indicate atoms that have a prediction, but for which no exact matches of the attributes are available in the BMRB. Grey boxes represent atoms that are not present in a given base.

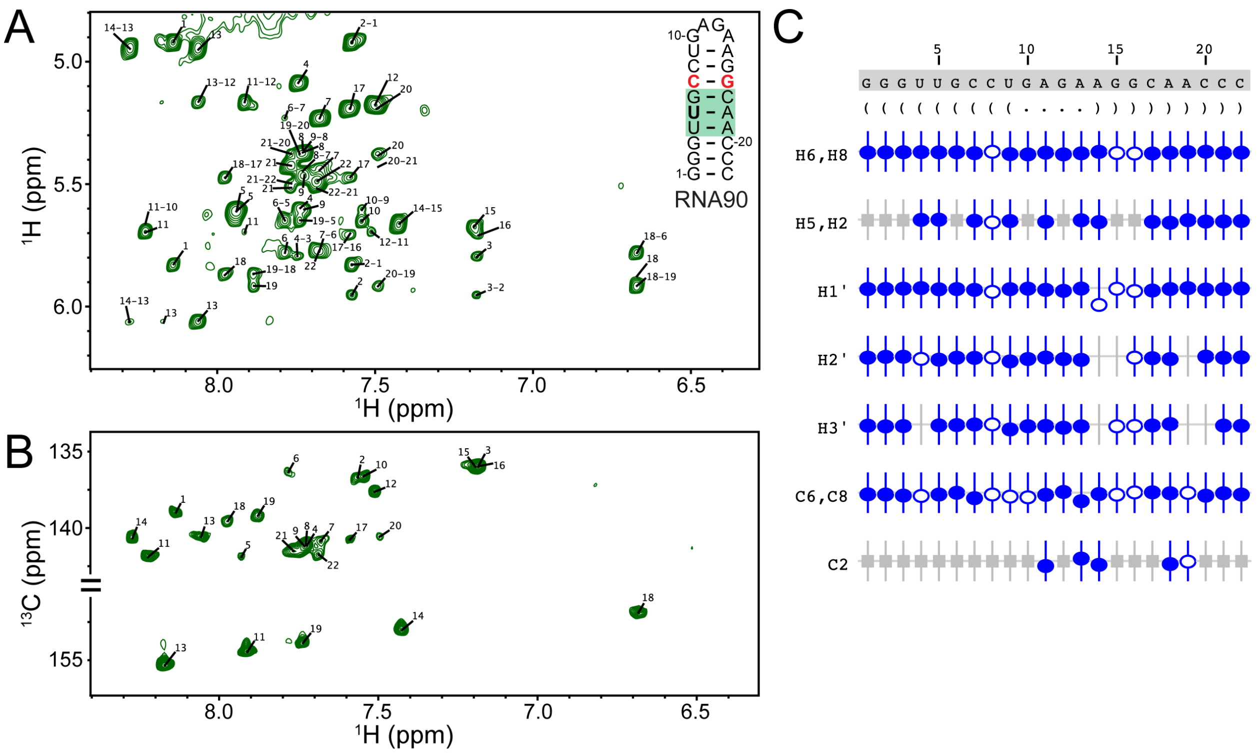


**Figure S7.** RNA90 chemical shift assignments. A) $^1$H-$^1$H NOESY, B) $^1$H-$^{13}$C HMQC, C) Summary of the sequence, secondary structure, and assignment validation for RNA90. The secondary structure is shown in Vienna format. NMRViewJ assignment summary to validate proton (H6/H8, H5/H2, H1′, H2′, H3′) and carbon (C6/C8, C2) assignments (bottom). Assigned chemical shifts for specific atoms are indicated with open and filled circles. The vertical offset of the circles indicates the deviation from the predicted values for that atom. Filled circles indicate that there are chemical shifts for atoms with the same set of attributes in the BMRB. Open circles indicate atoms that have a prediction, but for which no exact matches of the attributes are available in the BMRB. Grey boxes represent atoms that are not present in a given base.

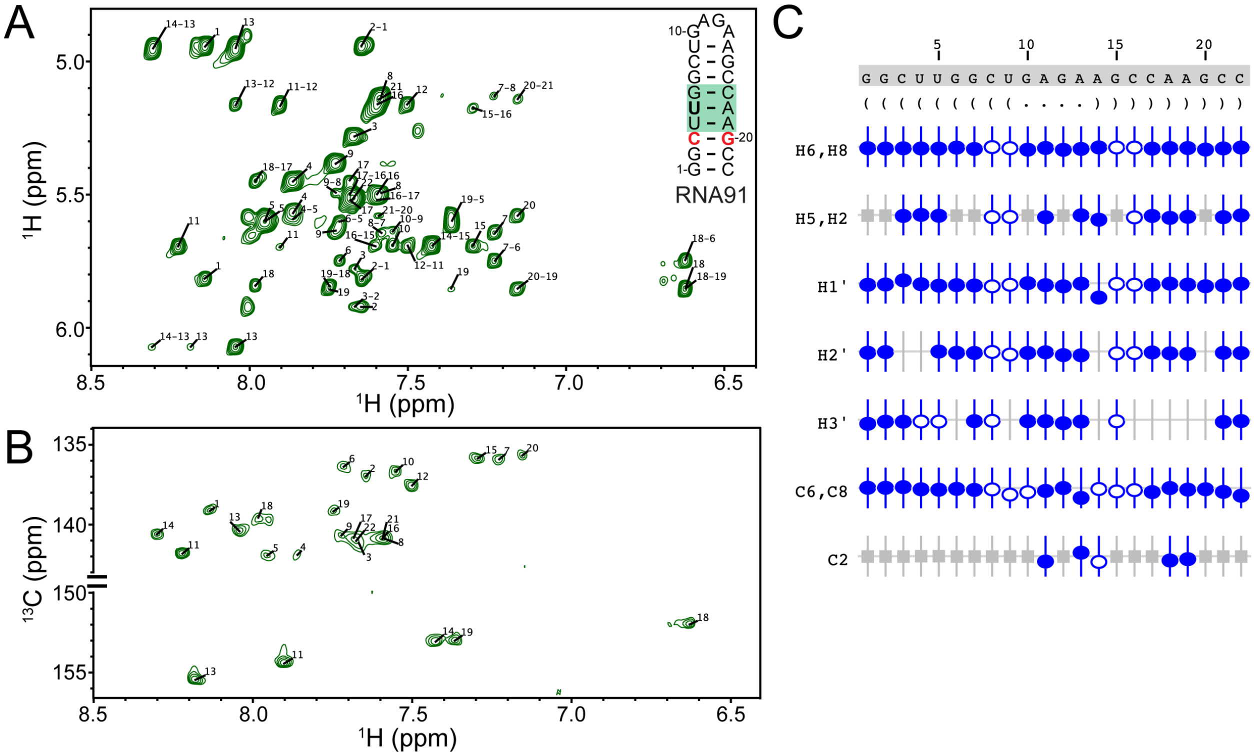

**Figure S8.** RNA91 chemical shift assignments. A) $^1$H-$^1$H NOESY, B) $^1$H-$^{13}$C HMQC, C) Summary of the sequence, secondary structure, and assignment validation for RNA91. The secondary structure is shown in Vienna format. NMRViewJ assignment summary to validate proton (H6/H8, H5/H2, H1′, H2′, H3′) and carbon (C6/C8, C2) assignments (bottom). Assigned chemical shifts for specific atoms are indicated with open and filled circles. The vertical offset of the circles indicates the deviation from the predicted values for that atom. Filled circles indicate that there are chemical shifts for atoms with the same set of attributes in the BMRB. Open circles indicate atoms that have a prediction, but for which no exact matches of the attributes are available in the BMRB. Grey boxes represent atoms that are not present in a given base.
